# A substrate recursion principle for biological information, with empirical anchoring through a templating-mode taxonomy

**DOI:** 10.64898/2026.06.23.734074

**Authors:** Boggavarapu Kiran

## Abstract

Biological inheritance can be treated as a class of catalytic templating reactions in which a daughter molecule, generated by a kinetic kernel acting on a parent template, is itself a substrate for the next round of the same catalysis. We give the physicochemical conditions under which such a reaction can support unbounded heritable molecular distinguishability. Four conditions on the template–operator pair organize the analysis: nonzero per-site information content under the activesite recognition kernel (R1), a count of independently variable recognized positions that grows without bound as the reaction extends (R2), catalytic closure under iteration, possibly through a reversible involution such as Watson–Crick complementation (R3), and stochastic drift of the kernel in its recognition alphabet (R4). A fifth, scope-defining condition (R5) restricts the principle to kernels that are intrinsic physicochemistry rather than externally optimized search. These conditions are necessary for two distinct outcomes, separable as two necessity results. The *capacity theorem* states that linear scaling of substrate Shannon capacity with reaction extent requires R1, R2, and R3 but not R4: a perfect copier transmits an exponentially large configurational ensemble while producing no novelty. The *generation theorem* states that diversification of the heritable configuration set beyond the deterministic closure of a finite initial repertoire additionally requires R4, because branching trajectories in the recognized alphabet are what produce innovation. Population-level kinetics follow as a corollary that organizes six attested templating reactions into a taxonomy, and as a finite-population, finite-horizon proposition tested over five inheritance kinetic schemes, in which only individual-level stochastic drift reaches the target within the model class. We test the framework on the recently characterized bacterial defense system Drt3b, which makes alternating poly(AC) DNA without using a nucleic acid template. The framework classifies Drt3b as a cyclic two-state catalytic templating channel with a 1-bit capacity ceiling, and predicts that the Glu26-to-Gln active-site mutant incorporates dG at 10% probability at the dA-selecting state; the published biochemistry reports 10.16%. Across 1,232 Drt3b homologs, the framework predicts and recovers a 15.7-fold elevation of dG misincorporation in six clades carrying the natural Glu26-to-Asp substitution at this gate. Substitutions at two universal gate residues, Arg253 (architectural) and Gly248 (selectivity), provide single-experiment site-directed mutagenesis tests of the framework’s predictions.

**POPULAR SUMMARY:** A bacterial defense protein called Drt3b, recently characterized in *E. coli*, synthesizes DNA with a strict alternating ACAC pattern without copying any template. Two conserved active-site residues, Glu26 and Arg253, are modeled as enforcing an alternating two-state catalytic cycle that selects which nucleotide enters at each step. This is sequence without a sequence template, and it does not fit the textbook picture of inheritance.

We treat inheritance as a class of catalytic templating reactions and ask which chemistries can support openended evolution. Two distinct requirements emerge. *Capacity*, the ability to transmit exponentially many distinct heritable configurations, requires three conditions on the template–catalyst pair: more than one monomer state recognized at each position, a position count that grows unboundedly with the reaction extent, and applicability of the catalysis to its own product. *Generation* of novel heritable configurations beyond what is already present requires a fourth condition: stochastic drift of the catalysis in its recognition alphabet. A perfect copier has capacity but cannot innovate; a drifting copier has both. Drt3b fails the second capacity condition, because its cycle has two states regardless of product length. The framework classifies six attested biological templating reactions as instances or partial instances of the same chemical specification, and it identifies two single-residue substitutions at Drt3b’s active site whose measured effects would test its predictions.

## I. INTRODUCTION

### A. The phenomenon and the puzzle

Biology runs on a small number of chemically distinct templating reactions for transmitting information across generations. Each is a catalyzed process in which a daughter molecule, generated from a parent template by the action of a catalyst, is itself the parent template for the next round of the same catalysis. DNA replication and ribosomal translation are the textbook cases: processive, ATP/GTP-coupled, polymeraseand ribosome-catalyzed monomer-by-monomer synthesis on a directing template. Prion propagation and amyloid templating add a second class, a conformational attractor propagated across protein molecules through nucleationand-elongation chemistry [1]. Modular biosynthetic assemblies (non-ribosomal peptide synthetases, polyketide synthases) compose a third: a spatial sequence of catalytic modules executes a fixed sequence of condensation reactions, with the product specified by the active-site chemistry of each module [2]. The eukaryotic ciliate cortex constitutes a fourth: cortical patterns of basal-body orientation are inherited at division through the action of the existing cortical lattice on newly assembled units, and grafted inverted patches propagate as inverted for hundreds of cell generations without genomic change [3, 4]. The recent characterization of bacterial defense-related templating systems, the DRT family [5–7], adds at least one more. Drt3b synthesizes alternating poly(AC) DNA without using a nucleic acid template, generating its product through a cyclic active-site catalytic mechanism rather than through Watson-Crick complementarity to an existing strand [5].

These reactions pose a physicochemical question, not a purely biological one: what conditions on a template–catalyst pair are necessary for it to support unbounded heritable molecular distinguishability under repeated catalytic action? The mechanisms have unrelated activesite chemistries, but their dynamical signatures separate cleanly. DNA replication transmits a configurational ensemble whose Shannon capacity grows linearly with sequence length. Prion propagation transmits a finite conformational free-energy minimum and accumulates nothing comparable. The field has no quantitative chemical-physics account of which reactions sit on which side of this line, or on what physicochemical basis the line is drawn. The question is surprisingly absent from the current literature, despite half a century of attempts to address adjacent ones.

### B. Prior work and the gap

The attempts have approached the substrate-level question from three directions. The information-theoretic line [8–10] treats copying-with-variation as a process whose stability depends on per-site error rate relative to genome length, generating the quasispecies framework and the error threshold *µ*^∗^ ≈ log *σ/L*. This work tells us how a substrate-borne replication process can fail dynamically, but not what makes a physical system a substrate at all. Szathmáry [10] introduced the distinction between limited heredity (a substrate with *K* states transmits a log_2_ |*A*| *K*-bit ceiling) and unlimited heredity (a sequence-based substrate transmits *L* log_2_ bits, scaling with length). The distinction is correct in spirit, but it is asserted as a typological observation rather than formalized as a structural condition on substrates.

A second line [11, 12] specifies abstract requirements on a replicator (faithful self-copying, heritable variation in fitness) without committing to any chemistry that realizes them. It is chemistry-blind: it does not say which physical realizations of the abstract requirements actually occur, or through what catalytic mechanism. The substrate-level line has attempted to specify the physical substrate directly. Schrödinger’s aperiodic crystal [13] was prescient but not formalized. The GARD-class compositional-inheritance models [14, 15] proposed that a compositional set of catalytic molecules transmits information across generations through preferential reproduction of similar compositions; Vasas *et al*. [16] showed, using GARD’s own machinery, that compositional inheritance fails the fidelity requirements for open-ended evolution. Pross [17] supplied a thermodynamic condition (dynamic kinetic stability) on replicators sustained far from equilibrium.

What is missing from all three lines is a single statement, at the chemical-physics level of the template–catalyst pair, of what conditions are necessary for a templating reaction to transmit unbounded heritable distinguishability. The information-theoretic work treats population dynamics on a given reaction, not the reaction itself; the replicator-criteria work is chemistry-blind; the substrate-level work has either gestured or been falsified. The Drt3b finding sharpens the gap. Drt3b is unambiguously biological, unambiguously templating, and unambiguously not a sequence-template polymerase. A framework that classifies it has to operate at the level of the catalytic mechanism itself.

### C. Here we show

We give such a framework. Four structural conditions on the template–catalyst pair, together with a fifth scope-defining condition, separate two necessity results: a capacity theorem and a generation theorem. Capacity is storage architecture under the catalytic kernel; generation is stochastic branching to neighboring configurations under the same kernel. Both are necessary for biology’s open-ended evolution, and they have different chemical requirements. A bounded-heredity corollary organizes six attested templating reactions into a taxonomy keyed to which condition each reaction fails, and a finite-population proposition characterizes the population dynamics each class hosts.

Section II states the framework: the formal definitions, the five conditions, the two theorems with proofs, the bounded-heredity corollary and its mode taxonomy, and the finite-population proposition. Section III reports the work that tests the framework: a descriptor-relative diagnostic apparatus, mode-specific scaling laws, quantitative anchoring on the Drt3 reverse-transcriptase system, a family-level analysis across 1,232 Drt3b homologs, sitedirected mutagenesis predictions, and a finite-population simulation, followed by the framework’s relationship to prior traditions, its limitations, and the orthogonal preconditions of life that lie outside its scope. Section IV states the conclusions. Materials and Methods give the formal setup, the simulation parameters, and the test protocols.

A note on vocabulary. We use *drift* rather than *error* or *variation* for the stochastic part of the catalytic kernel: the kernel produces what its physics produces, and there is no externally imposed target configuration the catalysis is trying to reach. We retain *error* for established field usage (Eigen’s threshold, and the per-position misincorporation rate *ε* defined against a Watson–Crick canonical product), where the technical meaning is fixed.

## II. DEFINITIONS, CONDITIONS, AND THEOREMS

We state the framework in three parts: definitions of heredity capacity and cumulative generation, the substrate recursion conditions, and two necessity theorems that separate capacity from generation. Each formal statement is preceded by a plain-language framing of what it claims and followed by a biological example.

### a. Heredity capacity and cumulative generation

Before specifying conditions on the substrate, we specify what the substrate is required to support. “Open-ended information generation” has been used loosely, and the looseness hides a quantifier ambiguity that determines which structural conditions are necessary. We separate two formal notions.

The first is *substrate heredity capacity* : how much information the substrate can transmit per recursion event, treated as an encoder property. The capacity is the supremum of mutual information over input ensembles supported on the recursively closed set ℛ_*L*_ := {*X* ∈ *X*_*L*_ : *O* ^*′*^ (*X*) ∈ *X*_*L*_ for all *t* ≥ 0, possibly via *ι*}:

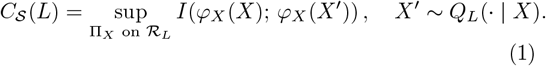

Restricting the supremum to ℛ_*L*_ makes R3 (catalytic closure under iteration) mandatory in the definition. A substrate failing R3, such as translation, where the protein product is not ribosome-readable as a next-round template, has ℛ_*L*_ = ∅ or trivially small, so *C*_*S*_(*L*) is zero or bounded by a constant. This distinguishes recursive heredity capacity from one-step channel capacity, which can be Ω(*L*) for non-recursive transfers (DNA → protein carries Ω(*L*) one-step bits but transmits them once and exits the substrate class). The substrate has *unlimited heredity capacity* if *C*_*S*_(*L*) = Ω(*L*); equivalently, in the discrete noiseless limit, if the set of heritably distinguishable and recursively readable configurations ℛ_*L*_ satisfies |ℛ_*L*_| ≥ 2^*cL*^ for some *c >* 0.

**Definition 1** (Heredity capacity). A substrate-operation pair (*X, O*) has unlimited heredity capacity if *C*_*S*_(*L*) = Ω(*L*) as *L* → ∞. Equivalently, in the discrete noiseless limit, |ℛ_*L*_| ≥ 2^*cL*^ for some *c >* 0.

**Remark 1**. The two formulations are coordinated through entropy: linear scaling of mutual information capacity (*C*_*S*_(*L*) = Ω(*L*) bits) is equivalent to exponential scaling of the recursively readable state count (|ℛ_*L*_| = Ω(2^*cL*^)). The first is the information-theoretic statement; the second is the combinatorial statement. They are the same fact at two levels of description.

The second is *cumulative generative recursion*: the substrate’s ability, starting from a finite initial repertoire *S*_0_ ⊊ *X*_*L*_ small compared to its reachable set, to produce heritable configurations *beyond the zero-drift closure of S*_0_ *under O*, through branching local alternatives in the same recognized alphabet. The qualifier “beyond the zero-drift closure” is essential. A deterministic operation can leave *S*_0_ (a counter chain 000 → 001 → 010 → … generates configurations outside any singleton *S*_0_), but the descendant set is then a single predetermined orbit. Cumulative generation in the chemical-kinetics sense requires that the future is not pre-specified: the kernel produces heritable branching alternatives in Σ at each step, so the descendant set is genuinely larger than the orbit closure under the catalysis’ canonical zero-drift output.

Formally, let *g*_*L*_(*X*) := *X*_ref_ (*X*) be the canonical zero-drift map of *O* (the kernel’s reference output: the Watson–Crick complement for a DNA polymerase, the exact alternating product for Drt3b). Let 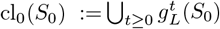 be the orbit closure of *S*_0_ under zero-drift iteration. Then (*X, O*) supports cumulative generation if the recursively readable descendant set reachable from *S*_0_ with positive probability under *Q*_*L*_ strictly contains cl_0_(*S*_0_) for all proper *S*_0_ ⊊ *X*_*L*_ with |*S*_0_| ≪ |ℛ_*L*_|, with the additional configurations differing from elements of cl_0_(*S*_0_) in local recognized-state positions (in the alphabet Σ). Using the canonical zero-drift skeleton avoids the pitfall of defining closure as a union over all deterministic restrictions of *Q*_*L*_: for stochastic *Q*_*L*_, such a union absorbs every positive-probability branch and trivializes the strict-containment condition. The zero-drift formulation picks one canonical deterministic skeleton, so R4-failure (zero drift, *Q*_*L*_ = *g*_*L*_) gives equality rather than impossibility.

**Definition 2** (Cumulative generative recursion). A substrate-operation pair (*X, O*) supports cumulative generative recursion if, for arbitrarily large *L* and any proper finite initial repertoire *S*_0_ ⊊ *X*_*L*_ with |*S*_0_| ≪ |ℛ_*L*_|, repeated application of *Q*_*L*_ generates a recursively readable descendant set whose heritable descriptor support strictly contains the zero-drift closure cl_0_(*S*_0_), with the additional configurations arising through branching local alternatives in the same recognized alphabet Σ as in (R1).

DNA illustrates both. The 4^*L*^-configuration space gives unlimited heredity capacity; the polymerase’s drift in the nucleotide alphabet gives cumulative generation. The two are conceptually independent. A hypothetical perfect copier of DNA satisfies Definition 1 (any of 4^*L*^ templates is faithfully preserved across iterations) but not Definition 2 (the descendant set from a finite *S*_0_ equals cl_0_(*S*_0_) itself, with no branching alternatives in Σ). A purely deterministic-counter substrate likewise fails Definition 2: the descendant set is bounded by the zero-drift orbit. Drt3b fails Definition 1 at the substrate level: at any product length, its two-phase cyclic dynamics distinguish only log_2_ 2 = 1 bit of phase, so *C*_*S*_(*L*) = *O*(1).

A note on what the kernel encodes. *Q*_*L*_(*X*^*′*^ | *X*) is the distribution over outcomes that the active-site freeenergy landscape, the available reactants, and the chemical driving (NTP hydrolysis or an analogous energy source) produce when the catalysis acts on *X*. The kernel does not encode a target product; every template in iterated *Q*_*L*_-application is the drifted product of prior runs of the same reaction, and no configuration is canonical at the kernel level.

### b. The substrate recursion conditions

The substrate-operation pair (*X, O*) is the formal object the principle constrains. *X*_*L*_ is the space of admissible configurations of size *L*; *ϕ*_*X*_ (*X*) ∈ Σ^*L*^ is the descriptor of *X* over the recognition alphabet Σ; *O* is the operation with kernel *Q*_*L*_(*X*^*′*^ | *X*) producing *X*^*′*^ = *O*(*X*), possibly composed with a reversible involution *ι* : *X*_*L*_ → *X*_*L*_. The involution covers the canonical case where the immediate product of *O* is the complement of the input (Watson-Crick complementarity produces 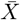, and the next round of replication on 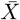 recovers *X*), allowing the recursion to close even when each individual *O* application produces a complement rather than a copy. Four conditions are structural and are what the necessity theorems require; the fifth is scope-defining.

**Definition 3** (Substrate recursion conditions). The substrate-operation pair (*X*, *O*) satisfies the substrate recursion conditions if:

#### (R1) Positive entropy per recognized site

At each addressable position *i*, the recognition step in *O* distinguishes at least two alternative monomer states drawn from a recognition alphabet Σ with |Σ| ≥ 2. The per-site entropy under the maximum-entropy distribution, *h*_site_ = log_2_ |Σ|, is positive.

#### (R2) Unbounded independently variable position count

The substrate has non-recurrent sites whose count *N* (*L*) grows unboundedly: lim inf_*L*→∞_ *N* (*L*) = ∞. “Independently variable” is meant in the firstorder, descriptor-relative sense: increasing *L* increases the count of heritable recognized degrees of freedom under the operation-relevant descriptor, and the substrate is not constrained to a finite recurrent phase repertoire. The condition does not require recognized states at distinct positions to be statistically independent in any strong joint-distribution sense; DNA polymerases exhibit context-dependent error rates and base-composition biases, and DNA still satisfies R2 because the count of independently variable positions grows with sequence length. Composition vectors, homogeneous concentrations, and bulk mixtures fail R2 because they lack addressable position. Cyclic systems with finite recurrent phase repertoires fail R2 because *N* (*L*) is bounded at the cycle length: product length can grow without bound while the count of heritable degrees of freedom stays at the cycle complexity *N*. This sharpening protects the Drt3b argument from a naive “product length grows” reading.

#### (R3) Same-operation recursion (with allowed involution)

The output *X*^*′*^ = *O*(*X*) is structurally an element of _*L*_ for which *O*(*X*^*′*^) is well-defined, possibly via a reversible involution *ι* such that *O*(*ι*(*X*^*′*^)) _*L*_. The recursion takes the form

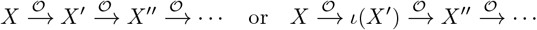

The condition excludes operations whose products are not readable by the same operation.

#### (R4) Heritable local drift in the recognized alphabet

The kernel *Q*_*L*_(*X*^*′*^ | *X*) has nonzero probability of producing configurations *X*^*′*^ ≠ *X*_ref_ (*X*) for any reference configuration, with the drift lying in the same alphabet Σ as the recognition step in (R1). Substrate content (the descriptor *ϕ*_*X*_ (*X*)) and substrate templating function (the role of *X* in *O* (*X*) → *X*^*′*^) are the same physical entity: drift in content is drift in the next iteration’s template.

#### (R5) Scope: non-teleological intrinsic operation

The kernel *Q*_*L*_ depends only on the physicochemistry of (*X, O*), not on an externally specified target configuration, loss function, or convergence criterion. (R5) is scope-defining rather than structural: it demarcates the domain in which (R1)–(R4) apply. Differential persistence of configurations may occur downstream over a finite population, but selection acts on products, not on the operation itself. Systems whose operation is externally optimized (gradient descent on a loss function, designed catalytic cycles with specified targets) are outside the principle’s scope.

The conditions in biological terms:

- **(R1)** requires the recognition step to distinguish at least two states at each site. DNA satisfies this: at each position the polymerase distinguishes A, T, G, and C, giving *h*_site_ = 2 bits. A polymerase that incorporates only T has *h*_site_ = 0 and fails (R1).
- **(R2)** requires the count of independently variable sites to grow without bound. DNA satisfies this; Drt3b’s cycle has only two phases regardless of product length, so its site count is bounded at 2 and (R2) fails.
- **(R3)** requires the operation to apply to its own output. DNA replication satisfies this through the involution 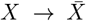. Ribosomal translation fails (R3): the protein product is not a substrate the ribosome can re-read, and recursion is recovered only through Mode 1 acting on the gene encoding the ribosomal machinery.
- **(R4)** requires drift in the alphabet the operation recognizes. DNA polymerases produce drift in the nucleotide alphabet (a ∼ 10^−9^ per-position rate against the Watson-Crick canonical product), and the drifted product is itself recognized at the next iteration. A perfect copier fails (R4); so does a system that drifts in an alphabet the polymerase does not read.
- **(R5)** requires intrinsic physicochemistry. DNA replication satisfies it; a neural network trained by gradient descent toward a target distribution does not, and is therefore outside the principle’s scope. (R5) does not say the externally optimized system fails biology; it says the principle’s structural analysis does not apply to it.

### c. Two necessity claims

The two definitions yield two distinct necessity claims. The first concerns heredity capacity, the substrate’s encoder properties as an information channel. The second concerns cumulative generation, its ability to innovate from a finite seed. Both are necessary for biology’s open-ended evolution, but they have different structural requirements, and the empirical phenomena are clearer when the two are kept separate.

**Theorem 1** (Capacity: necessity of R1, R2, R3 for unlimited heredity capacity). *Within the scope defined by* (R5), *if a substrate-operation pair* (*X*, *O*) *has unlimited heredity capacity in the sense of Definition 1, then it satisfies* (R1), (R2), *and* (R3).

**Theorem 2** (Generation: necessity of R4 for cumulative generative recursion). *Within the scope defined by* (R5), *if a substrate-operation pair* (*X*, *O*) *supports cumulative generative recursion in the sense of Definition 2 (its descendant set from finite S*_0_ *strictly contains* cl_0_(*S*_0_) *through branching local alternatives in* Σ*), then it satisfies* (R4) *in addition to* (R1), (R2), *and* (R3).

### d. Proof of Theorem 1 (sketch)

The argument is an upper-bound contrapositive. For any substrate-operation pair,

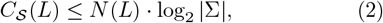

because 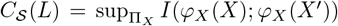 is bounded by the entropy of the descriptor, which has at most |Σ|^*N*(*L*)^ possible values. A matching lower bound requires a realizability condition: increasing *L* must actually create additional heritable recognized degrees of freedom under the operation-relevant descriptor. This is the operational content of R2, and it is what distinguishes a substrate with genuinely extensible recognized positions from one (a finite recurrent phase cycle) where product length grows but heritable degrees of freedom do not. Failure of (R1) gives |Σ| = 1, so log_2_ |Σ| = 0 and *C*_*S*_(*L*) = 0. Failure of (R2) gives *N* (*L*) ≤ *N*_max_ bounded, so *C*_*S*_(*L*) ≤ *N*_max_ log_2_ Σ, constant in *L* and therefore not Ω(*L*). Failure of (R3) means *O*^2^ is not defined within *X*_*L*_: the iteration terminates at the first application or transitions to a different substrate class on the second, so capacity inherited via a different upstream substrate counts as parasitic recursion rather than (*X*, *O*)’s own capacity. The contrapositive in each case gives the theorem. R4 is not invoked and is not necessary for capacity: a substrate satisfying R1, R2, R3 with *Q*_*L*_(*X*^*′*^ | *X*) = *δ*(*X*^*′*^ − *X*) (a perfect copier) has *C*_*S*_(*L*) = *N* (*L*) · log_2_ |Σ| and unlimited heredity capacity. This is why DNA without mutation would still be a high-capacity inheritance carrier: it would not innovate, but it could store and transmit.

### e. Proof of Theorem 2 (sketch)

R1, R2, and R3 are inherited from the requirement that the pair be coherent and recursively closed. Beyond these, suppose R4 fails. There are two sub-cases. (a) The kernel is deterministic with *Q*_*L*_(*X*^*′*^ | *X*) = *δ*(*X*^*′*^ − *g*_*L*_(*X*)), where *g*_*L*_(*X*) = *X*_ref_ (*X*) is the canonical zero-drift map. Then for any *S*_0_, the descendant set after *T* iterations is 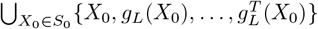, which equals cl_0_(*S*_0_). The deterministic operation may leave *S*_0_, but the descendant set is contained in the zero-drift orbit closure, with no branching alternatives, so strict containment fails. (b) The kernel produces drift in an alphabet Σ^*′*^ disjoint from the recognition alphabet Σ: the drifted configurations are not re-recognized at the next iteration, so they do not propagate as recursively readable descendants. Definition 2 requires the new configurations to arise through branching in Σ itself. In both sub-cases the substrate cannot produce a recursively readable descendant set strictly containing cl_0_(*S*_0_) through branching in Σ, contradicting Definition 2.

The “beyond deterministic closure” qualifier is the crucial logical move. An earlier formulation of the generation theorem said only that R4 is necessary for the descendant set to leave *S*_0_, which is false: a deterministic counter chain leaves any singleton *S*_0_ without satisfying R4. What R4 is actually necessary for is the descendant set being larger than the deterministic orbit closure, that is, the substrate producing genuine branching alternatives at each step rather than a single predetermined trajectory. The full proofs, including the argument that the conjunction’s scaling rate cannot be recovered from any subset of conditions, are in Supplementary Mathematical Appendix §5.

Theorem 1 explains why bounded-capacity substrates (Drt3b, prions) cannot support open-ended inheritance regardless of how much drift they produce: their structural ceiling caps the heritable repertoire at the wrong scaling. Theorem 2 explains why a high-capacity substrate without drift cannot innovate from a finite seed: it can preserve what exists but cannot accumulate novelty. Biology’s DNA-based heredity satisfies both: 4^*L*^ scaling gives capacity, and ∼ 10^−9^ per-position polymerase drift gives generation. The behaviors standardly described as natural selection, fitness, error thresholds, and evolutionary plateaus are downstream consequences, which we treat next as a corollary and a scoped proposition.

#### A. Bounded heredity and the substrate-mode taxonomy

A substrate failing one of (R1)–(R4) cannot reach the corresponding open-ended regime, but it is not thereby left with no heredity. What fails is the unboundedness of transmission (capacity, if R1, R2, or R3 fail) or the generative reach beyond a finite repertoire (if R4 fails). The heredity is bounded, and the bound is set by which condition fails.

**Corollary 1** (Bounded heredity). *If* (*X*, *O*) *is in the scope of (R5) and satisfies a strict subset of* (R1)–(R4), *then the substrate may exhibit heredity, but the heredity is bounded in a manner determined by which conditions fail. Substrates failing (R1) have configuration-space ceilings set by the per-site entropy (h*_site_ = 0 *at the limit, h*_site_ ≤ log_2_ *K for a K-state attractor); substrates failing (R2) have ceilings set by bounded position count N*_max_ *(cycle length, module count); substrates failing (R3) inherit parasitically through an upstream substrate that satisfies (R3); substrates failing (R4) exhibit fixed configurations or non-recursive drift without information growth*.

The corollary’s empirical instantiation is biology’s substrate-mode inventory. We document six modes, each a specific failure pattern of (R1)–(R4); capacity bounds and proofs are in Supplementary Mathematical Appendix §2–3.

##### Mode 1: complementarity-based 1D sequence templating

Substrate: a 1D linear sequence on a separate molecule. Mechanism: position-by-position chemical complementarity (Watson-Crick, *β*-sheet hydrogen bonding). Satisfies all four structural conditions within (R5) scope: *h*_site_ = log_2_ |*A*|*>* 0, unbounded position count *N* (*L*) = *L*, daughter strand a reactant for the next replication via complementarity (R3 via involution 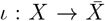), and heritable local drift through the polymerase kernel. Capacity: *C*_*S*_(*L*) = *L* log_2_|*A*| in the noise-free limit. Examples: DNA replication, Drt3a, retrons, telomerase, DRT9, *β*-sheet-mediated peptide replication.

##### Mode 2: code-encoded cross-descriptor templating

Substrate: a 1D sequence in *X* plus separable lookup-table machinery in. Fails (R3) at the substrate level: protein products are not readable by the ribosome as templates for further translation, so the recursion is recovered parasitically through the encoding gene, which is itself a Mode 1 substrate. Information transfer under uniform codon input is bounded by 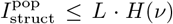, where *ν* is the codon-count distribution over output symbols (*H*(*ν*) = 4.218 bits/codon for the standard genetic code, including stop); the recursive capacity *C*_*S*_ itself is near zero because R3 fails. Example: ribosomal translation.

##### Mode 3: cyclic-structure-encoded templating

Substrate: a 3D protein structure with cyclic conformational dynamics having *N* distinct states. Fails (R2) in the sharpened sense: although product length can grow without bound, the count of independently variable recognized positions equals the cycle phase, bounded at *N* and recurrent (*N* (*L*) = *N* regardless of product length). The macro-level capacity is consequently bounded at log_2_ *N*, and the output is periodic with period *N*. This distinguishes the framework’s R2 from a naive “product length grows” reading: Drt3b produces arbitrarily long products, but its heritable distinguishable information stays at log_2_ *N* bits. Examples: Drt3b producing (AC)_*n*_ via *N* = 2 conformational states [5]. DRT7 is the boundary case at *N* = 1, where log_2_ 1 = 0: there is no positional information capacity, only structural channel selectivity, so we classify DRT7 as a Mode 3 boundary rather than a positional-templating instance.

##### Mode 4: conformer state-templating

Substrate: a discrete conformational state of a protein. Fails (R1) at the macro level (the recognition alphabet is a finite attractor repertoire of *K* conformers) and fails (R2) trivially (a single effective position, the conformer assignment, rather than an extensible series). Capacity: *C*_*S*_(*L*) ≤ log_2_ *K*, length-independent. Examples: prion propagation, amyloid self-conversion, DRT1 filamentous oligomer [6]. Prion strain repertoires confirm that some drift is transmitted across iterations, but the transmissible drift lies in a small attractor set rather than an extensible alphabet.

##### Mode 5: modular conveyor templating

Substrate: a spatial sequence of *N* structurally distinct modules within a single multi-modular assembly. Fails (R2) at the module count (position count is bounded by *N*, not extensible without adding modules) and (R3) at the substrate level (peptide products are not the assembly line; recursion is parasitic on Mode 1 acting on the gene cluster). Capacity: *C*_*S*_(*L*) = *N* · log_2_ *k* for *L* ≥ *N*, with output length bounded by *N*. Examples: non-ribosomal peptide synthetases, polyketide synthases.

##### Mode 6: 2D surface-position templating

Substrate: a spatial register on a 2D pattern, where the existing pattern biases the assembly of new units that do not themselves store the pattern. Mode 6 has extensible geometric positions (R2 holds: surface position count grows with area) and per-site entropy in the unit-type alphabet (R1 holds with multiple unit types). The failures are at (R3) and (R4): the 2D pattern does not autonomously self-copy with heritable local drift independent of genome-encoded parts. Pattern copying is not a same-operation recursion of the surface; it depends on Mode 1 acting on the genome to produce the constituent proteins, and pattern drift independent of the genome’s drift cannot accumulate as heritable variation in the recursive sense. The biological instance is the eukaryotic ciliate cortex [3, 4]: cortical patterns of ciliary basal-body orientation are inherited at division through templating action of the existing cortex on newly assembled units, and microsurgically grafted inverted patches propagate as inverted for hundreds of generations without genomic change. The framework retrodicts the cortex as a real but bounded inheritance system, parasitic on Mode 1 in the same structural sense as Mode 2.

The taxonomy is an *empirical inventory* of currently analyzed biological templating architectures, not a deductive exhaustion of all possible substrates satisfying or partially satisfying (R1)–(R4). Every characterized DRT system fits one of Modes 1, 3, 4, plus the framework’s out-of-scope boundary (Tables II–III); Modes 2, 5, 6 are anchored elsewhere (translation, NRPS/PKS, ciliate cortex). The inventory is provisional in the sense any empirical taxonomy is: new biology may force additions, as Drt3b forced Mode 3 to be made explicit. A more principled future taxonomy might derive modes from substrate dimensionality, recognition cardinality, and recursion topology as orthogonal axes within the principle.

#### B. Finite-population proposition

The structural conditions classify substrates; the population dynamics those substrates host follow as a scoped, finite-horizon claim. An earlier formulation stated a strong universal claim, that R3or R4-failing mechanisms are structurally capped at mechanism-specific fitness ceilings under any population dynamics. That claim is too strong: a wholesale-redraw mechanism with full support over the configuration space can in principle sample the target by chance given infinite time, even though the probability is negligible in any finite regime. The issue is efficiency, not absolute reachability, so we state the result as a proposition scoped to the tested finite-population landscape model and framed as an adaptive-efficiency argument. The proposition is tested by simulation in Sec. III F.

**Proposition 1** (Finite-horizon plateau under nonlocal or absent drift). *In a finite population of size K, evolving under selection sharpness β toward a target phenotype on the tested-class landscape (single-target Hamming, Ldimensional alphabet, static), over a horizon T <* ∞:

1. *Mechanisms whose substrate-operation pair fails (R4) at the individual level (exact-copy systems) cannot exceed the maximum fitness present in their founding lineage repertoire. They are bounded by initial-population properties, not by their own variation capacity*.
2. *Mechanisms with nonlocal (R4) (wholesale redraw at rate r >* 0*) sample novelty with full support but do not accumulate local improvements; under the tested update rules, the probability of reaching F*_max_ *within horizon T is dominated by random hitting time on the full configuration space and remains negligible for L »* 1.
3. *Mechanisms satisfying (R3) and (R4) with local drift at the individual level are not structurally barred from cumulative climbing toward F*_max_. *Whether climbing actually occurs depends on landscape topology, finite-population drift, mutation load, neutral-network connectivity, and other conditions outside the principle’s scope*.

In words: a population of perfect copiers cannot improve past the maximum fitness already present in its founding generation, since selection sorts what exists but generates no new variants; a population of wholesale-redraw copiers can sample anywhere but accumulates local improvements inefficiently, because each redraw discards lineage information; only individual-level local drift converts variation into cumulative climb. The positive direction is bounded: it says only that local-drift mechanisms are not structurally barred from climbing, not that they always climb.

## III. RESULTS AND DISCUSSION

### A. A descriptor-relative apparatus distinguishes templating from biasing participants

The framework’s empirical anchoring requires an instrument that classifies templating events by mode. We define a templating event as the tuple (*X, O, S*, Δ*G*; *ϕ*_*X*_, *ϕ*_*Y*_) → *Y*, where *X* is the template, *O* the operator, *S* the substrate pool, Δ*G* the free-energy driving, and *ϕ*_*X*_, *ϕ*_*Y*_ the descriptor maps. The diagnostic apparatus is a joint observable in three coordinates.

The first is *population mutual information*,

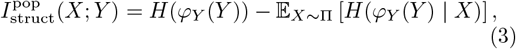

the expectation taken over a stated template population *X* with distribution Π. It is bounded above by *H*(Π) and is identically zero when *X* is degenerate, so it measures *realized* information transfer in a stated population, distinct from substrate capacity.

The second is *channel mutual information* against a stated comparison ensemble *C* of templating channels,

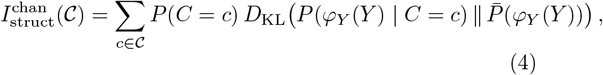

with mixture distribution 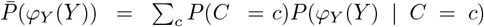. Unlike 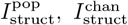 does not vanish for fixed-template channels: a channel with a single fixed template still produces a distinguishable output distribution against alternative channels.

The third is *per-base mechanism fidelity*, resolved into two components when the channel has cyclic structure. *Phase fidelity f*_phase_ is the fidelity of cycle-state alternation; *monomer fidelity f*_monomer_ is the fidelity of correct monomer choice within each state. The triple 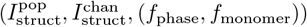 is the apparatus signature, and mode classification uses the joint signature rather than any single coordinate.

We distinguish realized population MI from *substrate capacity* 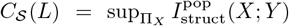, a property of the substrate class rather than of any specific population. *C*_*S*_ is what enters the principle’s (R1) condition; 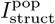 is what the apparatus measures on a particular biological system. A claim that *X* is a structurally specific template requires two controls. The bulk-matched control *X*_bulk_ produces an output *Y*_bulk_ with *k*th-order matched marginals but no position-specific information; the structure-scrambled control *Y*_scram_ has the same composition as *Y* with positional pattern shuffled. Real templating must give 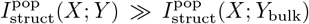 and 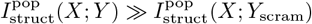 to a stated significance threshold.

We validate the apparatus on the canonical case. A simulated Watson-Crick replication system with template length *L*, alphabet size 4, and per-position misincorporation rate *ε* should give 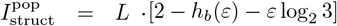. Across 30 sweep cells, the empirical 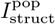 matches theory to within 0.6% (Figure 1A). The bulk-matched control discriminates cleanly: across 24 non-uniform sweep cells, separation ratios between genuine templating and bulk-matched control range 450*×* to 1350*×* (Figure 1B). The signature is robust to the choice of comparison ensemble for 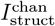: across 10 random four-channel ensembles drawn from a 10-channel pool, every channel’s mode classification was stable.

**FIG. 1:**
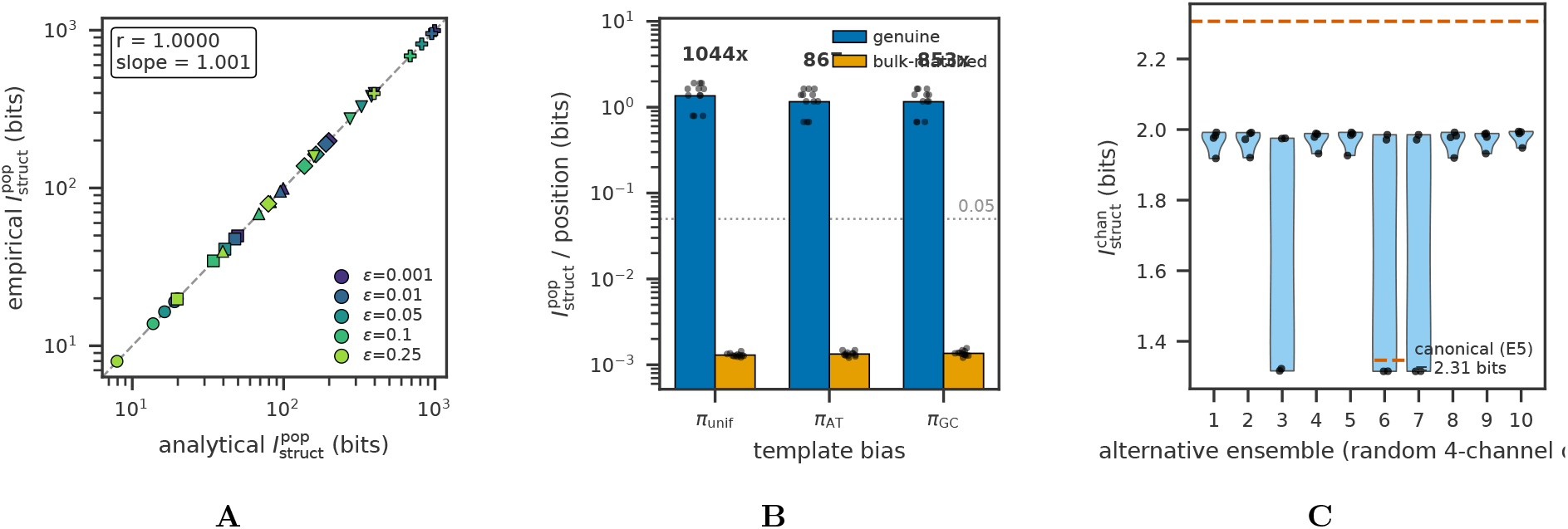
Apparatus validation and bulk-matched control. (**A**) Empirical 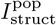 versus analytical prediction across the *ε × L* sweep, showing close tracking of analytical capacity in noise-free Mode 1 templating (test A1). (**B**) Separation ratio 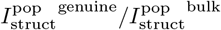 across three template-bias conditions, showing *>* 20*×* separation between genuine templating and bulk-matched controls (test A2). (**C**) Comparison-ensemble robustness: 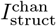 across 11 alternative comparison ensembles (test G3); the qualitative classification of Drt3a as Mode 1 and Drt3b as Mode 3 is preserved across all ensembles, confirming that the apparatus’s classificatory verdict is stable under reasonable analyst choices.

### B. Mode-specific scaling laws

We tested Corollary 1’s mode-specific capacity predictions directly. A two-state cyclic active site, sweeping *N* ∈ {2, 3, 4, 5, 6, 8, 10} with low-noise channel (*ε* = 10^−3^), should approach 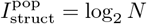 from below. Empirical measurements approach the bound to within estimator precision: 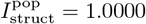 bits at *N* = 2, 2.3219 bits at *N* = 5, 3.3219 bits at *N* = 10 (Figure 2A). For each *N*, 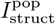 is constant across *L* from *L* = *N* to *L* = 50*N* to within 0.2% (worst case at *N* = 2, *L* = 2; the bound is *<* 0.01% for *N* ≥ 3). Mode 3 information saturates while Mode 1 information scales linearly with *L* (Figure 2B), confirming the extensibility-failure prediction within finite-sample noise.

**FIG. 2:**
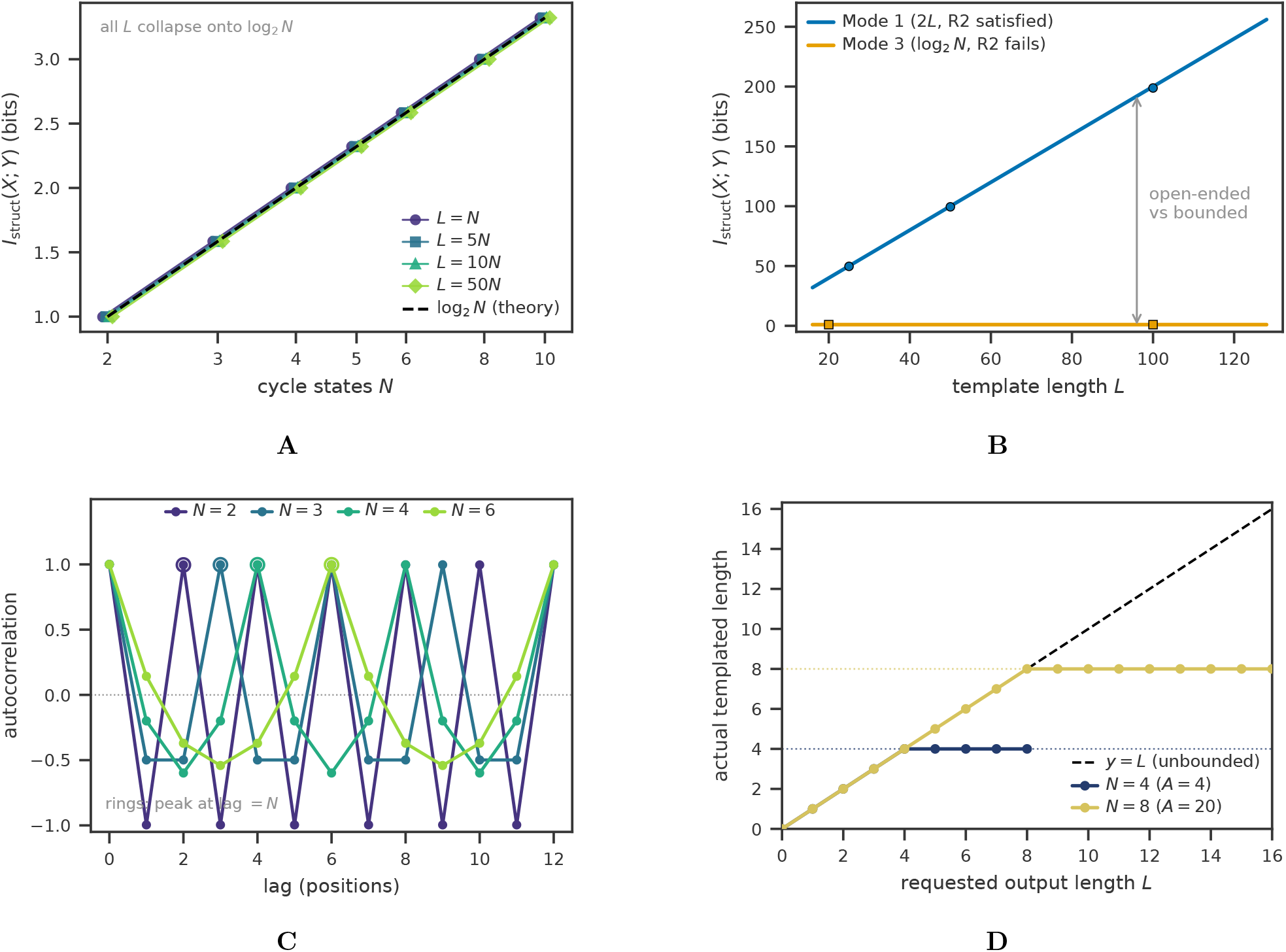
Mode-specific information-capacity scaling. (**A**) Mode 3 information capacity 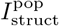 saturates at log_2_ *N* as cycle states *N* increase (test B), regardless of template length *L*, confirming that the macro-level capacity bound is set by the cycle’s recurrent phase repertoire and not by output length. (**B**) Mode 1 capacity scales linearly in *L* (slope log_2_|*A*| = 2) while Mode 3 saturates at log_2_ 2 = 1 bit; the structural difference between (R2)-satisfying and (R2)-failing substrates is visible in this single panel. (**C**) Mode 3 output autocorrelation peaks at lag = *N* for *N* ∈ {2, 3, 4, 6}, the canonical signature of cyclic-phase templating. (**D**) Mode 5 output length is capped at the module count *N* regardless of requested *L* (test C), confirming the (R2)-failure-at-module-count diagnosis.

The autocorrelation of Mode 3 output peaks at integer multiples of *N*, with peak value matching the noise-model prediction (1 − *ε*)^2^ + *ε*^2^*/*(*N* − 1) (Figure 2C). A Mode 5 simulation sweeping module count *N* ∈ {2, 4, 8, 16, 32} at alphabet sizes 4 and 20 confirms linear scaling in *N*, with a sharp length limit: positions beyond the module count carry zero information about the template (Figure 2D). For Mode 2, we simulated the standard genetic code applied to uniformly random DNA templates of *L* codons. The empirical 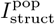 matches the analytical value *L* · *H*(*ν*) = 4.218*L* bits within finite-sample noise (4.214 at *L* = 64, *n* = 5000). The Mode 1-vs-Mode 2 ratio at matched DNA length is 6*/*4.218 ≈ 1.42. A nested two-level evolution simulation, with selection on phenotype and the codon-to-amino-acid lookup table itself encoded as a Mode 1 substrate, produced the predicted dynamic: 500 generations drove population fitness from 5% to 43%, with code fidelity converging to 52% as selection acted on phenotype while the recursion acted on the gene encoding the code. The simulation illustrates the mechanism by which Mode 2 inherits parasitically through Mode 1; it does not by itself confirm the biological claim about translation, which rests on independently established molecular biology.

### C. Quantitative anchoring to the Drt3 system

The Drt3 system [5] provides a direct biological test. The complex contains two reverse transcriptases and a noncoding RNA. Drt3a uses a conserved ACACAC region of the ncRNA as a Watson-Crick template (Mode 1) to produce poly(GT) DNA. Drt3b synthesizes the complementary poly(AC) strand without any nucleic acid template, using a cyclic active site in which two conserved gating residues impose alternating state-specific nucleotide selectivity: Glu26 gatekeeps dA selection through hydrogen-bonding to the N^6^ amine of dA, and Arg253 stabilizes dA through a cation-*π* interaction while making three hydrogen bonds with the Watson-Crick edge of dC at the C-state. The Glu26-to-Gln point mutation perturbs dA gating selectively, and the in vitro mutant produces alternating poly(AC) with approximately 80% dA / 20% dG misincorporation at the dA-selecting position while preserving dC selectivity [5, Fig. 4J]. The same protein fold appears in AbiK with different activesite residues, where the system produces random ssDNA [5]. We model Drt3b as a cyclic active-site templating channel; whether the gating residues undergo discrete physical flipping in the structural-biology sense is a separate question that the apparatus-level classification does not require.

We classify these systems by parameterizing Markovchain channels from the published biochemistry and asking whether the apparatus recovers the expected classifications. Throughout, we label each claim as observed (from Deng *et al*. [5]), derived (analytical consequence), simulated (Markov-chain apparatus on parameterized channels), or predicted (untested experimental target); Table IV maps every claim to its category.

Drt3b WT, parameterized from the (AC) alternating product, gives 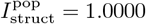 bits, saturating the cyclic-channel capacity bound log_2_ 2 = 1 bit. Phase fidelity *f*_phase_ ≈ 0.99 (cycle alternation intact) and monomer fidelity *f*_monomer_ ≈ 0.99 at both states. Periodicity peaks at lag 2 with peak autocorrelation 0.9802; output marginal frequencies are 0.497 dA, 0.497 dC, 0.003 dG, 0.003 dT; the separation ratio against the bulk-matched control ranges 234*×* to 1031*×*.

Drt3b Glu26-to-Gln, parameterized with the paper’s 80%/20% dA/dG split at the dA-selecting state, gives 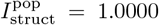 bits (capacity bound preserved) and 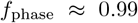 (cycle architecture preserved), but *f*_monomer_ ≈ 0.80 at the A-state with C-state selectivity unchanged. The marginal G fraction rises to 0.102, matching the analytical prediction 0.5 *×* 0.20 = 0.10 for half-cycle dG misincorporation (Deng et al. report 0.1016); the periodicity peak drops to 0.83; the separation ratio against the bulk-matched control falls to ∼ 100*×* (Figure 3A,B). The apparatus classifies the mutant as a cyclic channel with degraded monomer selectivity at one state but intact phase architecture, a distinction that motivates the architectural-versus-selectivity gate classification below.

**FIG. 3:**
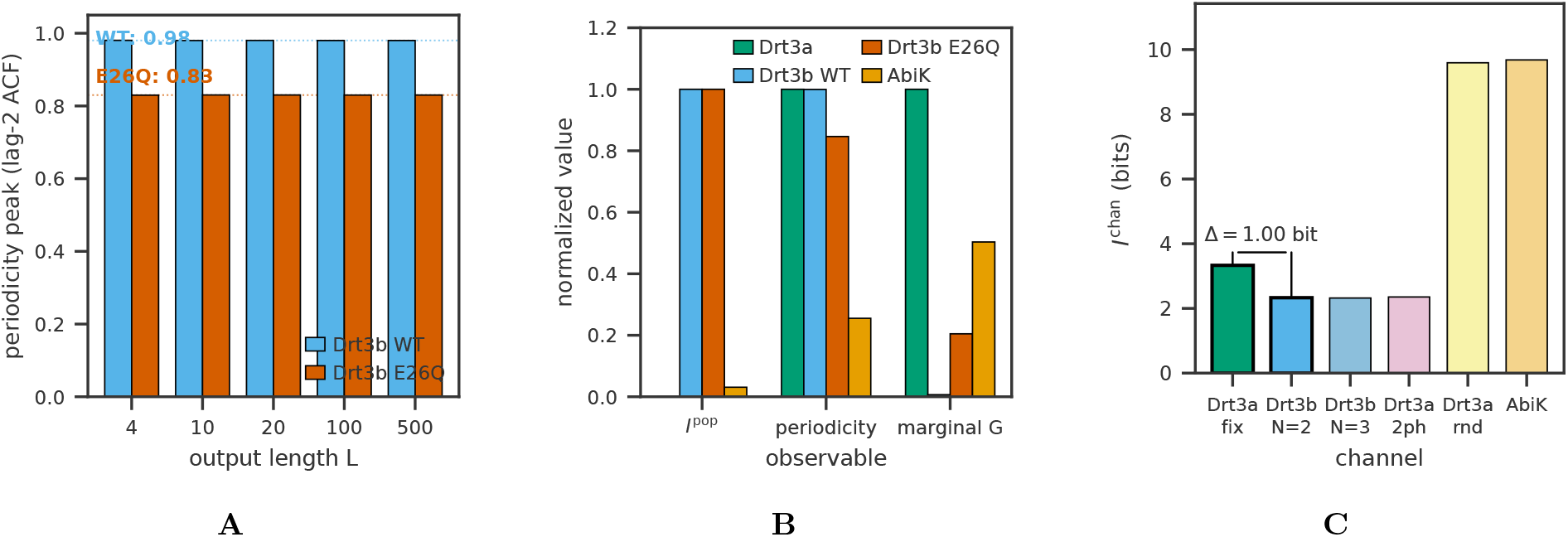
Apparatus signature on the Drt3 system. (**A**) Drt3b WT versus Glu26-to-Gln periodicity peaks (test E2): WT shows *f*_phase_ ≈ 0.99, the mutant shows *f*_phase_ ≈ 0.83, confirming the analytical prediction of half-cycle dG misincorporation in the gate-broken mutant (0.5 *×* 0.20 = 0.10 marginal G fraction; observed 0.1016 in Deng et al. 2026 Fig. 4J). (**B**) Apparatus triple (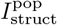, periodicity peak, marginal G fraction) separating WT Drt3b, Glu26-to-Gln, AbiK, and Drt3a in joint-observable space (test G2); the four systems classify cleanly despite producing nearly identical alternating output for some pairs. (**C**) Channel-mutual-information separation: against the stated five-channel comparison ensemble, the Drt3a Watson-Crick channel gives 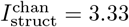 bits and the Drt3b *N* = 2 channel gives 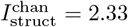 bits, yielding 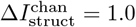 bit between Mode 1 and Mode 3.

**FIG. 4:**
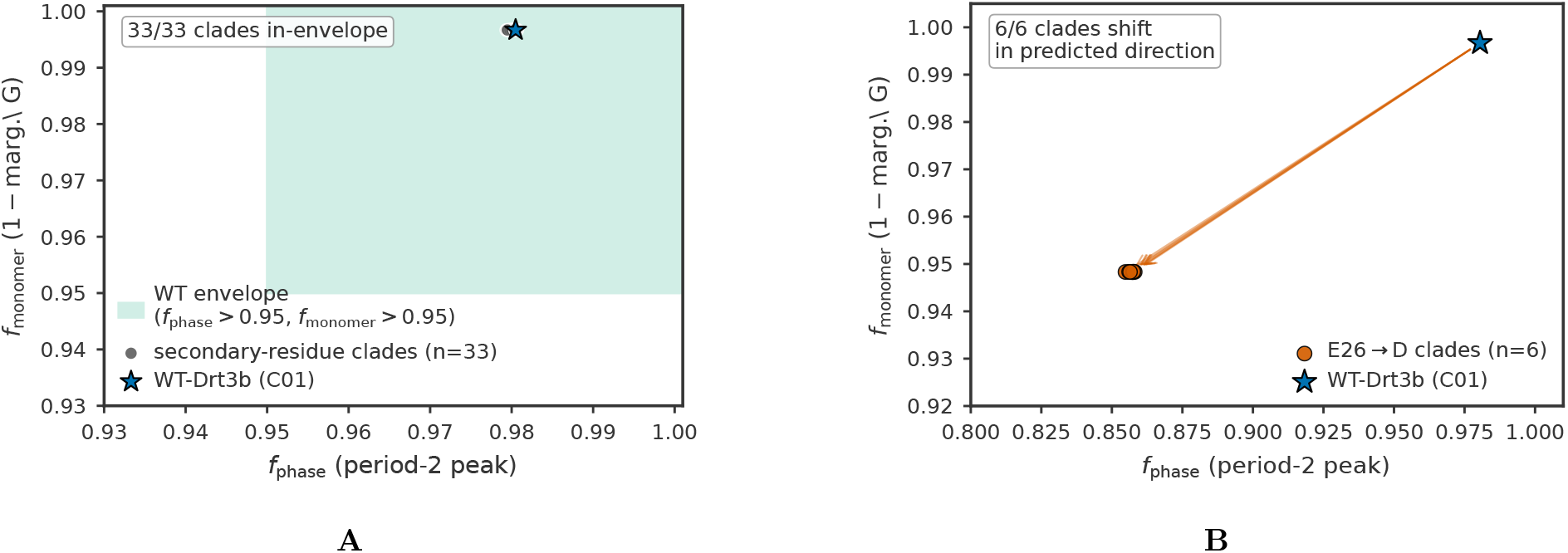
Drt3b family-level apparatus application across 1,232 homologs. (**A**) In-envelope behavior of 33 secondary-residue clades (test F): all 33 fall within the WT-Drt3b apparatus envelope, supporting the inference that secondary-residue diversity does not break the cyclic-conformational mode. (**B**) Glu26-to-Asp selectivity-gate signature in 6 primary-gate clades (C06, C14, C20, C27, C28, C42): the marginal G fraction rises from ∼ 0.003 in WT-like clades to ∼ 0.052 in the Glu26-to-Asp clades, a 15.7-fold elevation, consistent with residual hydrogen-bonding flexibility from the shortened Asp side chain. The periodicity peak drops to ∼ 0.857 as the arithmetic consequence of A-state selectivity loss (analytical 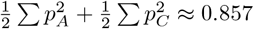, not as an independent phase failure: cycle architecture is intact (Arg253 invariant in all 6 clades). All 6 clades show the framework-derived selectivity shift, supporting the classification of Glu26 as a primary selectivity gate distinct from the architectural gate Arg253. Apparatus results are computed on Markov channels parameterized from residue assignments at conserved positions, not from raw product sequencing.

AbiK same-fold non-templating, parameterized from the published random-product description, gives 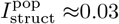 bits and *f*_monomer_ ≈ 0.25 (random over four nucleotides). The apparatus correctly excludes AbiK from the cyclic-templating class and from any structurally specific template class.

Drt3a (Mode 1) and Drt3b (Mode 3), treated as distinct channels in a five-channel comparison ensemble, separate by 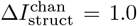 bit despite producing nearly identical alternating output (Figure 3C). The channel-as*X* observable distinguishes mechanism even when output statistics match: Drt3a’s *f*_monomer_ characterizes Watson-Crick fidelity against an external ACACAC template, while Drt3b’s *f*_phase_ and *f*_monomer_ characterize the cyclic active-site channel. The numerical separation is ensemble-dependent, but the joint signature’s qualitative discrimination is stable across ensemble choices, as the robustness check above shows. WT Drt3b and the Glu26to-Gln mutant both saturate 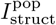 at the 1-bit bound, yet differ sharply in marginal G fraction (0.003 versus 0.102) and periodicity peak (0.98 versus 0.83). The mutant and AbiK both produce non-WT compositions but differ in 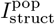 (1.0 versus 0.03) and *f*_monomer_ (0.99 versus 0.25). Collapsing the joint signature to a single score discards the information that distinguishes the systems.

### D. Family-level application across 1,232 homologs

We applied the apparatus across 1,232 Drt3b homologs identified from the Deng et al. supplementary alignment. Each homolog was assigned to a clade by primary-gate residue (Glu26, Gln26, Asp26, other) and secondary-residue identity at six conserved positions identified through structural alignment. The residue partition follows the Deng paper’s structural assignments: Arg253 (universally invariant) is the architectural gate, enforcing cycle phase via cation-*π* stabilization of the templating dC; Gly248 (essentially invariant) is a selectivity gate at the C-state via steric exclusion of dG; Glu26 is a primary selectivity gate at the A-state, mimicking a templating nucleobase via two hydrogen bonds with the N^6^ amine of incoming dA [5]. The apparatus makes two predictions. Clades with WT-like primary and secondary residues should fall in-envelope at the WT-Drt3b signature (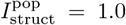 bit, marginal G fraction *<* 0.005). Clades with the Glu26-to-Asp primary-gate substitution shorten the carboxylate side chain and degrade A-state selectivity, so the framework predicts elevated dG misincorporation at the A-state with the cycle architecture preserved (Arg253 is invariant in all six such clades).

Of 33 secondary-residue clades assayed, all 33 fall in-envelope (Figure 4A). Of the 6 Glu26-to-Asp primary-gate clades (C06, C14, C20, C27, C28, C42), all 6 show the predicted shift: the marginal G fraction rises from ∼ 0.003 in WT-like clades to ∼ 0.052, a 15.7-fold elevation in dG misincorporation at the A-state, consistent with the residual hydrogen-bonding flexibility allowed by the shortened Asp side chain (Figure 4B). The periodicity peak drops from ∼ 0.98 to ∼ 0.857, the arithmetic consequence of A-state selectivity loss rather than an independent architectural failure. With state-A residency probabilities (0.85, 0.10, 0.025, 0.025) over (A, G, C, T) and state-C unchanged at 0.99 for C, the analytical period2 self-match probability is 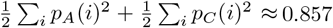, matching the simulation to three decimal places.

The cycle architecture is intact in all six clades (Arg253 invariant), and the apparatus localizes the failure to the A-state selectivity channel rather than the cycle’s phase structure. The application is consistent with the cyclic-templating classification holding across the Drt3b family rather than only at the reference sequence; it is an apparatus-on-parameterized-channels result, not a direct measurement of homolog products, which Deng et al. do not deposit.

### E. Universal-gate site-directed-mutagenesis predictions

The architectural-versus-selectivity distinction translates into structurally distinct site-directed mutagenesis predictions, with the family-level Glu26-to-Asp data already providing an empirical anchor on the selectivity side. Arg253 is an architectural gate: its substitution disrupts the cation-*π* stabilization that holds the cycle in its alternating phase, predicted to collapse 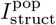 to near zero with the periodicity peak dropping to ∼ 0.43. Gly248 is a selectivity gate at the C-state: its substitution alters monomer selectivity within an intact cycle, predicted to preserve 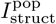 near WT while elevating marginal G to ∼ 0.15. Glu26 is a selectivity gate at the A-state, structurally analogous to Gly248 but on the complementary half-cycle; the family-level Glu26-to-Asp data confirm the selectivity-gate signature empirically (marginal G ∼ 0.052, cycle phase preserved). Arg253-toAla is therefore the cleanest single-experiment test the framework offers, with 100*×* predicted separation from WT 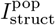.

Arg253-to-Ala is predicted to collapse 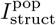 to 0.0090 bits (from WT 0.9997 bits), with the marginal G fraction rising to 0.1265 and the periodicity peak dropping to 0.4325; the cycle architecture is destroyed and the system reduces to compositional bias only. An observed 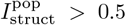 bits would refute the claim that Arg253 is the architectural gate. Gly248-to-Ala is predicted to preserve 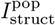 at 0.9966 bits, with the marginal G fraction rising to 0.1516 and the periodicity peak dropping to 0.7469; the cycle stays intact and the dG misincorporation comes from a selectivity shift at one state. An observed 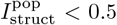 bits would make Gly248 architectural and the gate-class distinction would fail. Predictions for partial substitutions (Arg253-to-Lys, Arg253to-His, Gly248-to-Val, Gly248-to-Asp) and the double mutant (Arg253-to-Ala + Gly248-to-Ala) are stated in Table V. The double mutant is the strongest combined test of the gate-class ordering: under the framework’s prediction the architectural collapse should dominate the joint signature, whereas a dominant selectivity signature would show the ordering to be wrong.

### F. Finite-population dynamics: bounded-heredity plateaus

We tested Proposition 1 in a pre-specified five-mechanism characterization^1^ spanning the inheritance landscape from M0 (no recursion: fails R3 and R4) to M4 (canonical Mode 1: satisfies R1–R4 within R5 scope). M0 is stateless, with phenotype freshly drawn each generation. M1 is lineage-level fixation: at lineage founding a random template is drawn, all lineage members share it, and reproduction copies the lineage label with no individual-level drift (satisfies R3 at the lineage level, fails R4). M2 is lineage-level fixation with re-draw rate *r* ∈ [0, 1] (R3 at the lineage level, partial R4 via wholesale re-draw rather than local drift). M3 is individual-level exact copy without drift (R3 at the individual level, fails R4). M4 is individual-level copy with local drift (R3 and R4 at the individual level).

Plateau heights at generation 1000, averaged across 30 replicates with target length *L* = 32, *K* = 400 agents, and selection sharpness *β* = 10: M0 = 0.249 (chance), M1 = 0.472, M3 = 0.498, M2 at *r* = 0.10 = 0.533, M4 = 0.972 (Figure 5A; Table VI). Only M4 reaches near-target fitness in the tested model class. M1 and M3 plateau at the maximum fitness present in the initial population’s lineage repertoire, the (R4) failure producing no new drift. M2 has a regime-invariant sweet spot at *r* = 0.10: too little re-draw and the system cannot escape sub-optimal initial draws, too much and the lineage advantage washes out. The sweet spot is invariant within 0.05 across *β* ∈ {2, 5, 10, 20} and *L* ∈ {16, 32, 64, 128}.

**FIG. 5:**
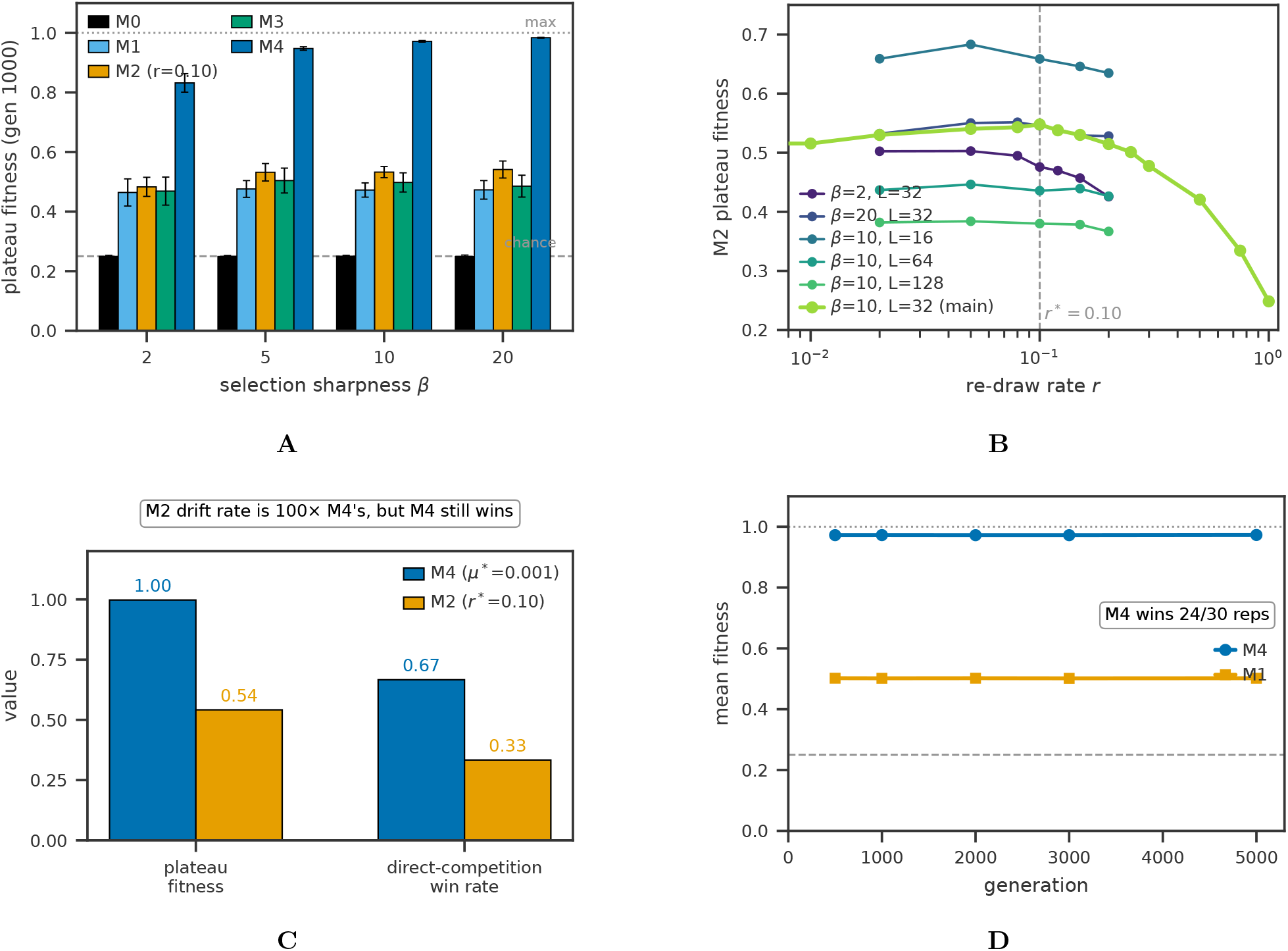
Population-dynamic plateaus from substrate condition failure. (**A**) Mechanism-specific plateau heights 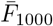 across selection sharpness *β* ∈ {2, 5, 10, 20} (test H3): M0 (R3, R4 fail) plateaus at chance (0.250); M1, M3 (R4 fails) plateau near initial-population maximum (∼0.47 to 0.50); M2 at *r* = 0.10 (partial R4) plateaus at ∼ 0.53; M4 (R1–R4 satisfied) reaches ∼ 0.97. (**B**) M2 sweet-spot at *r*^∗^ ≈ 0.10, regime-invariant within *±*0.05 across *β* ∈ {2, 5, 10, 20} and *L* ∈ {16, 32, 64, 128} (test H5). (**C**) M4 versus M2 head-to-head at matched per-offspring drift rate (test H5): M2 introduces 2.40 new positions/offspring at *r*^∗^ = 0.10; M4 introduces 0.02 at *µ*^∗^ = 0.001; M4 still wins 67%/33% in direct competition, demonstrating that targeted local drift converts to fitness gain more efficiently than wholesale lineage re-draw. (**D**) Long-horizon (5000 generations) M4 versus M1 trajectories at equal-start initial conditions (*K*_*M*1_ = *K*_*M*4_ = 200): M4 wins 24/30 replicates; M1’s plateau is durable but bounded.

A direct head-to-head test confirms the proposition’s content: lineage-level re-draws and individual-level local drift are not interchangeable input sources in the tested model. M2 at *r*^∗^ = 0.10 introduces 2.40 expected new positions per offspring; M4 at *µ*^∗^ = 0.001 introduces 0.02. M2’s drift rate is 100*×* M4’s, yet M2 reaches plateau 0.548 while M4 reaches 0.997, and in direct competition at balanced start M4 wins 67%, M2 wins 33%, with no ties. Targeted local drift converts input into fitness gain more efficiently than wholesale lineage re-draw in the tested landscape: the structural distinction matters, and rate alone does not. The qualitative claim survives at long horizons. At 5000 generations with equal-start *K*_*M*1_ = *K*_*M*4_ = 200, M4 wins outright in 24/30 replicates against M1, the remaining 6/30 reflecting cases where M1 fixes early before M4’s drift-and-selection cycle produces a winning lineage; the limited-heredity plateau is durable under the tested update rules, not a transient. At more asymmetric initial conditions (*K*_*M*1_ = 80, *K*_*M*4_ = 320 or *K*_*M*1_ = 20, *K*_*M*4_ = 380), M4 wins 30/30.

The proposition’s prediction for the eukaryotic ciliate cortex is consistent with observation (Figure 6). The cortex is built each generation from genome-encoded proteins (re-draw at every cell division) while the existing pattern biases the assembly (lineage-level fixation), operationally close to M2 with low re-draw rate, exactly the regime where the framework predicts limited but heritable variation. Cortical inheritance is documented to propagate non-genomic structural variation for hundreds of generations [3], consistent with M2’s plateau persistence; the cortex does not provide an independently extensible sequence-like substrate, consistent with M2’s bounded ceiling.

**FIG. 6:**
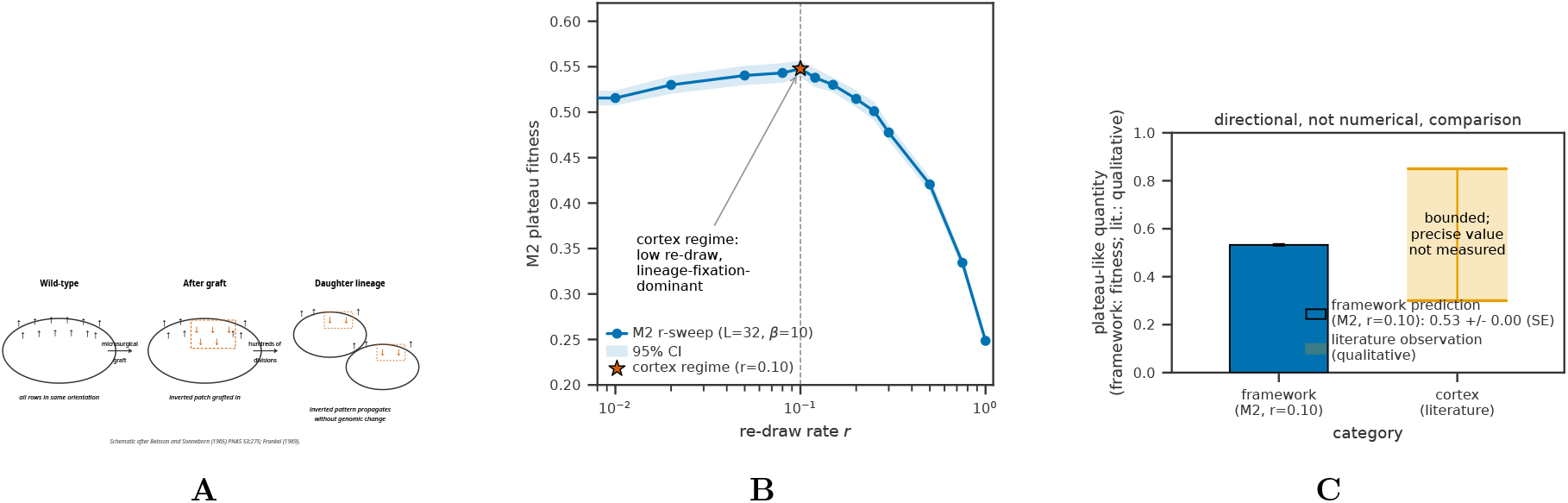
Mode 6 reframing and ciliate cortex bounded-heredity retrodiction. (**A**) Schematic of cortical inheritance after Beisson and Sonneborn [3]: a microsurgically grafted patch of inverted ciliary basal-body rows propagates as inverted across hundreds of cell divisions in *Paramecium* without genomic change, demonstrating that the cortex inherits non-genomic structural variation. (**B**) Operational analog within the M2 mechanism class: at every cell division, cortical units are re-drawn from genome-encoded parts, and the existing pattern biases the orientation of newly assembled units. The framework places the cortex in the M2 regime with *r* ≈ 0.10 (low re-draw, lineage-fixation-dominant), consistent with the Beisson-Sonneborn observation. (**C**) Predicted versus observed: the framework predicts bounded but persistent heredity for Mode 6 substrates (M2 plateau at ∼0.53 in simulation), and the literature reports bounded but persistent cortical inheritance (hundreds of generations of pattern propagation, no qualitatively new cortical structures accumulated over evolutionary timescales). The comparison is directional, not quantitative: the literature’s bounded heredity is a qualitative observation, not a fitness measurement.

### G. What the framework establishes

The framework gives biology’s templating chemistries a structural specification they have lacked. The four conditions (R1)–(R4) are properties of the template–catalyst pair under the scope of (R5), not of the population-level dynamics the catalysis hosts. Capacity and generation are separable structural requirements: capacity (R1, R2, R3) bounds the Shannon information the reaction can transmit per round, and generation (R4) controls whether iterated runs can branch off the deterministic closure of a finite initial repertoire. Keeping them separate clarifies which biological chemistries fail what. Drt3b (Mode 3) fails capacity at R2, its cycle having bounded independently variable positions, so drift status is moot once capacity is capped. Prions (Mode 4) fail capacity at both R1 and R2. Translation (Mode 2) recovers capacity parasitically through Mode 1 because its own product fails R3. Population-level kinetics (selection, fitness landscapes, error catastrophe, evolutionary plateaus, lineage formation) are downstream of which structural conditions the underlying chemistry satisfies. Re-grounding the analysis at the catalytic-reaction level reframes the question “what supports biology?” as “what templating chemistries can host biology’s recursion?”

The structural conditions are unambiguous; their operationalization is descriptor-relative. The conditions are stated at the kernel level, but applying them to a particular reaction requires a descriptor choice (*ϕ*_*X*_, *ϕ*_*Y*_) that fixes what counts as a recognized position and what counts as a distinguishable monomer, and the apparatus inherits this descriptor-relativity. This is not a weakness: a descriptor-free taxonomy would be either trivially true (all biology is templated) or untestable (no apparatus could be defined). The framework’s commitment is that, given an admissible descriptor choice, the conditions generate stable mode classifications across reasonable descriptor variations, which we have demonstrated for Drt3b across descriptor choices spanning the natural range.

The framework’s relationship to natural selection is one of layered description. Selection is real as a statement of which catalytic kernels persist in finite populations under environmental filtering, but it is not a force acting inside the kernel itself. The reaction runs because the active-site free-energy landscape, the reactant pool, and the chemical driving allow it to run; certain configurations are produced because the kernel produces them; certain configurations fail to carry forward because they are not produced, or are produced unstably. Selection-talk is descriptive vocabulary applied to the population-level distribution of catalytic kernels, a deflationary reading of selection’s role inside the reaction rather than a denial of its population-level reality.

### H. Relation to prior traditions

The framework is complementary to the autopoieticclosure tradition. Rosen [18], Maturana and Varela [19], Pattee [20], and Hofmeyr [21] ask how a living system produces its own templates, the catalysts that read those templates, and the metabolic cycles that produce both. The present framework specifies which classes of templating chemistry can serve as the inheritance carrier in such a self-producing system, identifying Modes 1 and 2 as the only attested candidates. We do not address the origin or self-maintenance of the catalytic machinery itself.

The strongest empirical challenge in the prior literature is yeast prion biology [22, 23]. Prion conformations are real, ecologically prevalent, and evolutionarily active in wild yeast populations, generating heritable phenotypic variation that selection acts on. The framework’s response is not that prion inheritance does not exist, but that its inheritance carrier is parasitic on Mode 1: what is transmitted prion-to-prion is a conformational attractor on a polypeptide whose primary sequence is genome-encoded. The prion can stabilize or accelerate adaptation by exposing cryptic genetic variation, but the carrier of heritable distinguishability is the gene encoding the prion-prone protein, itself a Mode 1 substrate. Prions are bounded conformational templating chemistries (Mode 4) supplementing an underlying Mode 1 reaction.

The strongest formal challenge is collectively autocatalytic sets of the GARD/compositional-inheritance class [14, 15]. Vasas *et al*. [16] showed, using the GARD kinetics directly, that compositional inheritance fails the fidelity requirements for open-ended evolution: highfidelity transmission of composition across cell divisions requires a lognormal distribution of catalytic rate constants so narrow as to be biologically unreachable. The present framework reaches the same verdict by a different route. GARD-class compositional inheritance fails capacity-theorem R2, because composition vectors lack independently variable positions that scale with reaction extent, and fails generation-theorem R4, because fluctuation in composition does not propagate as drift in a recognized alphabet under a per-site catalytic kernel (there is no per-site recognition step). The two arguments are independent: Vasas et al.’s is a dynamical-kinetic-fidelity result for a specific kinetic scheme, ours a structural argument from the catalytic-kernel geometry. They reach the same conclusion, and the present framework supplies the chemical-physics reason behind the empirical failure.

Mode 6 (the ciliate cortex) is the framework’s positive prediction for a 2D-surface templating chemistry. The cortex has been documented since 1965 as a non-genomic structural inheritance system, with grafted patches of inverted basal-body rows propagating as inverted for hundreds of cell generations without genomic change [3, 4]. The framework places it as a bounded-heredity supplementary chemistry dependent on Mode 1: the cortex satisfies R1 and R2 at the surface level (the lattice has multiple unit-types at independently variable positions) but fails R3 and R4 (the surface does not autonomously self-catalyze its own copying with stochastic drift independent of the genome). Pattern transmission depends on Mode 1 acting on the genome to produce the cortical proteins, and pattern drift independent of genome drift cannot accumulate as heritable variation in the recursive sense.

### I. Limitations and scope

The biological inventory of six modes is provisional, and new biology may force additions. The taxonomy is descriptive: the modes are derived from attested examples rather than from a deductive enumeration of all possible chemistries satisfying or partially satisfying (R1)–(R4). A more principled future taxonomy would derive modes from active-site dimensionality, recognition cardinality, and recursion topology as orthogonal axes within the framework. The predictions have been tested in coarse-grained Markov-chain models of the catalytic kernel; an atomistic free-energy calculation of Drt3b’s active-site cycle would test whether the two-state picture survives realistic energy-landscape geometry. The DRT family classifications rest on published mechanism descriptions rather than direct apparatus application to raw product reads, and access to deposited Drt3-product sequencing would close that gap.

### J. Orthogonal preconditions of life as we know it

The framework specifies the necessary structural conditions on a template–catalyst pair for it to support open-ended Darwinian inheritance. This is one of several essential preconditions for life as we know it, not the only one. Metabolism, the coupled catalytic network that maintains the cell as a far-from-equilibrium system, is orthogonal: a population of inheritance-capable substrates without metabolic closure is a chemistry experiment, not a living lineage. Compartmentalization, the boundary that keeps a lineage’s chemistry coherent against environmental dilution, is orthogonal: an inheritance reaction in bulk solution disperses its descendants into the medium. Sustained chemical driving (NTP hydrolysis, redox gradients, photochemical input) is orthogonal: without driving, the catalytic kernel runs once and stops, and (R3) and (R4) collapse to detailed-balance equilibria. Each has its own theoretical literature: Pross [17], Maturana and Varela [19], Hofmeyr [21] for closure-tradition treatments of metabolism, the protocell and vesicle-encapsulation literature for compartmentalization, and nonequilibrium statistical mechanics for chemical driving. Among the substrate-level conditions for the informational spine of biological inheritance, however, the framework is, to our knowledge, the one not previously stated in unified form. Prior work supplied individual conditions or pairs (Schrödinger’s aperiodic crystal, Eigen’s error threshold, Wagner’s neutral networks, Hull’s replicator ontology, Hofmeyr’s (*F, A*)systems, Vasas et al.’s GARD refutation; Supplementary §5.8 traces the correspondence), but the capacity/generation separation under a single (R5) scope is new.

## IV. CONCLUSIONS

Biology’s templating chemistries sort into a small inventory keyed to the chemical character of the catalytic kernel. The conditions that separate open-ended from bounded inheritance are properties of the template–catalyst pair, and they split into a capacity requirement and a generation requirement with different chemical content: a perfect copier has capacity without generation, and a drifting copier has both. DNA replication is the only attested chemistry that meets every condition. Translation, the modular biosyntheses, and the ciliate cortex inherit parasitically through DNA because their own products are not re-read; prions and Drt3b are bounded at the substrate level by a small conformational alphabet and a two-state cycle, respectively.

The framework anchors quantitatively to the Drt3 reverse-transcriptase system. It predicts 10% marginal dG misincorporation for the Glu26-to-Gln mutant against a published 10.16%, and it predicts and recovers a 15.7-fold dG elevation across six natural Glu26-to-Asp clades among 1,232 homologs. Two single-residue substitutions at universal gate residues would test its central claim about gate-class architecture: Arg253-toAla should collapse the cyclic architecture, and Gly248to-Ala should produce selectivity loss without architectural collapse. A result discordant with either prediction would refute that claim. Metabolism, compartmentalization, and sustained chemical driving remain orthogonal preconditions for life with their own theoretical lineages. Whether the structural specification holds across novel chemistries is an empirical question: the family analysis, the placement of seven additional DRT systems, the Mode 6 reframing, and the population-dynamics characterization make the principle a bet that future templating-chemistry findings will decide, rather than a stipulation.

## V. MATERIALS AND METHODS

### A. Mathematical setup

A templating event is the tuple (*X, O, S*, Δ*G*; *ϕ*_*X*_, *ϕ*_*Y*_) → *Y* where *X* is the template, is the operator (separable molecular machinery, possibly empty or coincident with *X*), *S* is the substrate pool, Δ*G* is the free-energy budget, and *ϕ*_*X*_, *ϕ*_*Y*_ are descriptor maps on template and product spaces. When *ϕ*_*X*_ = *ϕ*_*Y*_ the templating is same-descriptor; otherwise it is cross-descriptor and *O* implements the coupling.

The substrate recursion conditions (R1)–(R5), Definitions 1–2, Theorems 1–2, Corollary 1, and Proposition 1 are formalized in Supplementary Mathematical Appendix §5. The diagnostic apparatus is the triple 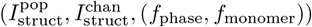 defined formally in Supplementary Mathematical Appendix §1. We summarize the operational definitions used in the main paper. Population mutual information 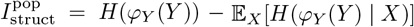 is estimated via plug-in entropy estimators on empirical joint distributions. Channel mutual information 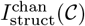 is estimated as the mixture-MI under a stated comparison ensemble using *n*_samples_ ≥ 5000 per channel. Per-base fidelities *f*_phase_ (cycle-state alternation) and *f*_monomer_ (canonical-monomer selection within a state) are estimated as empirical correctness rates against canonical-output functions. The substrate capacity 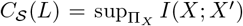 is computed analytically per mode (Appendix §2) rather than estimated.

The bulk-matched control *X*_bulk_, *Y*_bulk_ preserves *k*thorder statistics of *Y*. For *k* = 1, *Y*_bulk_ is drawn iid from the empirical first-order composition. For *k* = 2, *Y*_bulk_ is drawn from the maximum-entropy first-order Markov chain preserving empirical nearest-neighbor frequencies. The structure-scrambled control *Y*_scram_ has the same composition as *Y* with positional pattern shuffled.

### B. Simulation framework

Simulations were implemented in Python 3.10 using NumPy 1.24 and Matplotlib 3.7. Random seeds were set explicitly for reproducibility (per-replicate seed = 42 + cell idx *×* 100 + rep idx). Computations ran on a 64-CPU compute node. Code and raw simulation outputs are available at 10.5281/zenodo.20272479 (full submission-state snapshot); see Data Availability for the relationship to the earlier pre-specification deposit at 10.5281/zenodo.20060972.

### C. Test A.1: Mode 1 length-scaling validation

Templates *X* of length *L* over alphabet {*A, C, G, T*} drawn uniformly. Products *Y* generated by Watson-Crick channel with per-position misincorporation rate *ε*. Sweep *ε* ∈ {0.001, 0.01, 0.05, 0.10, 0.25} and *L* ∈ {10, 25, 50, 100, 200, 500} with *n* = 5000 per cell. Per-position MI *I*(*X*_*i*_; *Y*_*i*_) computed by plug-in estimator. Theoretical comparison: 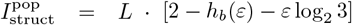.

### D. Test A.2: bulk-matched control discrimination

Three template biases tested (*π*_uniform_, *π*_AT−skew_, *π*_GC−skew_). Bulk control draws *Y*_bulk_ iid from analytical output marginal *Q* at each position, with *X*_bulk_ uncorrelated. Pass: 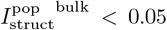 bits, separation ratio *>* 20*×*.

### E. Test B: Mode 3 information saturation

Two-state cyclic active site simulation. Sweep *N* ∈ {2, 3, 4, 5, 6, 8, 10}, *L* ∈ {*N*, 5*N*, 10*N*, 50*N*}, *ε* = 10^−3^, *n* = 5000 per cell. 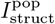 computed against template ensemble equal to phase index. Theoretical bound: 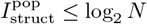.

### F. Test C: Mode 5 module-bounded scaling

Modular conveyor channels with *N* modules, alphabet sizes |*A*| ∈ {4, 20}. Sweep *N* ∈ {2, 4, 8, 16, 32}, *L* = 5*N*, *n* = 5000 per cell. Theoretical bound: 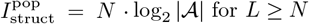 for *L* ≥ *N*.

### G. Test D: Mode 2 codon-degeneracy capacity

Standard genetic code (NCBI table 1). Random DNA templates length *L* codons, *L* ∈ {16, 32, 64, 128}, *n* = 5000 per cell. Output measured per amino acid (with stop). Theoretical value: *L* · *H*(*ν*) = 4.218*L* bits.

**TABLE 1:**
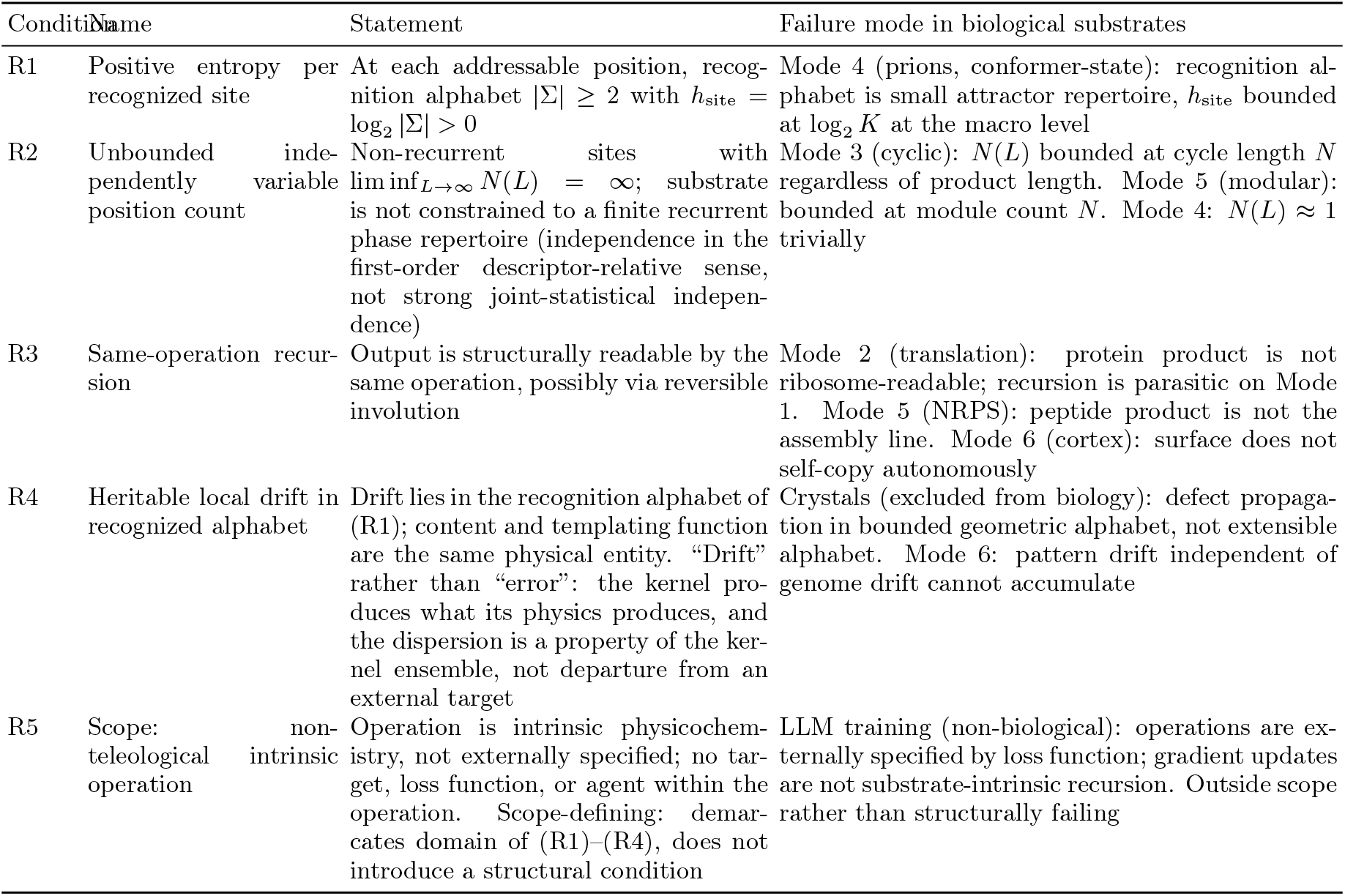
Substrate recursion principle: four structural conditions plus one scope-defining condition.

**TABLE 2:**
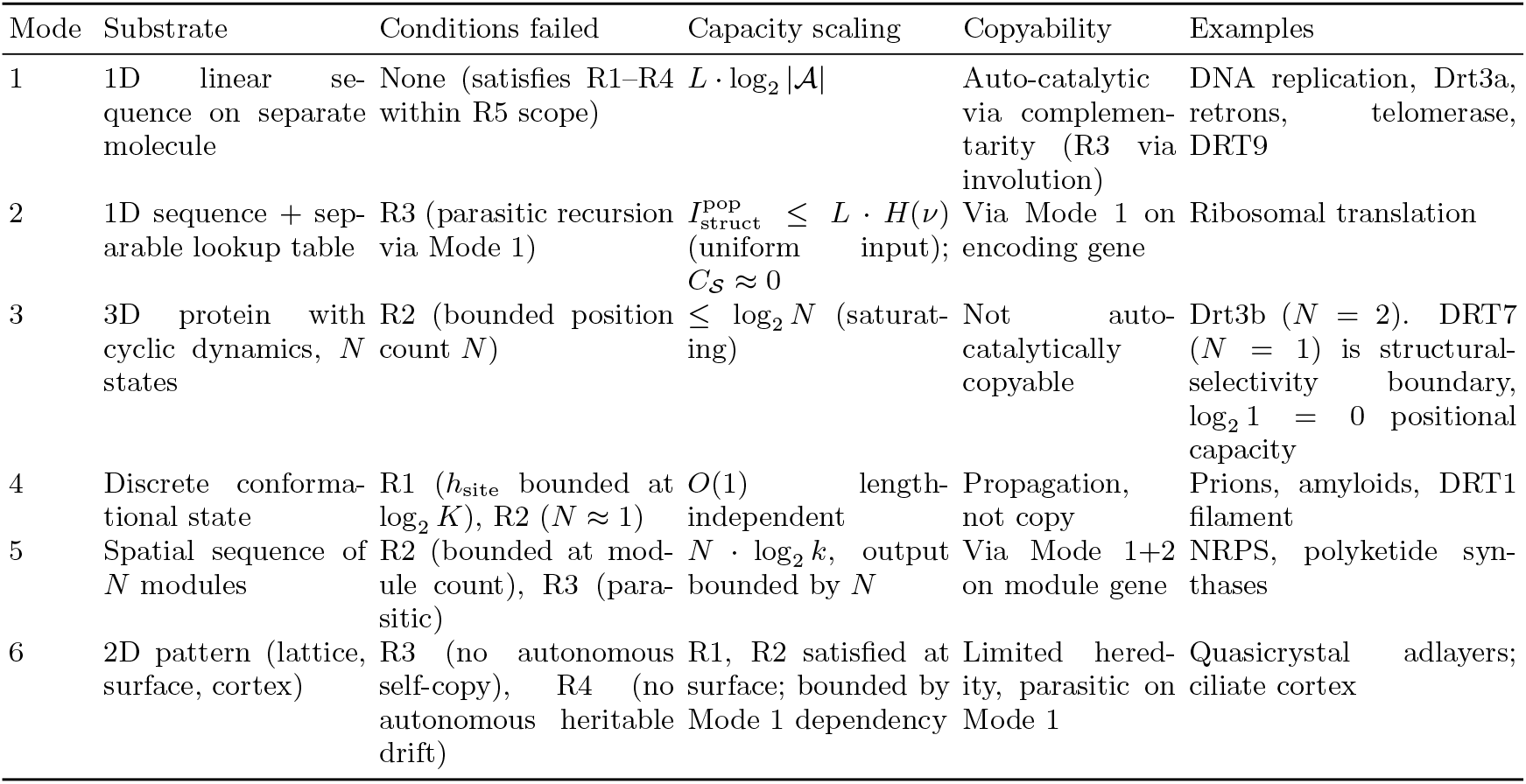
Six modes of biological templating, classified by substrate recursion principle conditions. The Mode 2 entry gives the one-step channel mutual information under uniform codon input; its recursive capacity *C*_*S*_ is near zero because R3 fails.

**TABLE 3:** DRT family classification under the framework.

| System | Mode | Mechanism summary | Reference |
| --- | --- | --- | --- |
| DRT1 | Mode 4 | Filamentous quaternary state with ssDNA effector | [6] |
| DRT2 | Mode 1 | Rolling-circle reverse transcription on ncRNA template, *neo* effector | [24, 25] |
| DRT3a | Mode 1 | Watson-Crick on ACACAC ncRNA region, produces poly(GT) | [5] |
| DRT3b | Mode 3 ($N = 2$) | Cyclic protein-conformational gating, produces poly(AC) | [5] |
| DRT4 | Out of scope | Random ssDNA, biasing-participant (toxin) | [26] |
| DRT6 | Out of scope | Random ssDNA, structural homolog of DRT4 | — |
| DRT7 | Mode 3 boundary ($N = 1$) | Single-state arginine-rich pocket, $\log_2 1 = 0$ positional capacity; structural channel selectivity remains, but no positional templating; classified as boundary case rather than full Mode 3 instance | [7] |
| DRT9 | Mode 1 | Poly-U ncRNA template, produces poly(dA) homopolymer | [27] |
| DRT10 | Mode 1 | Translocation-based reuse of short ncRNA template; telomerase ancestor | [28] |

**TABLE 4:** Epistemic status of the Drt3 claims. Results in this paper fall into four categories. *Observed* = published experimental fact from Deng *et al*. [5]. *Derived* = arithmetic or analytical consequence of an observed fact under the framework. *Simulated* = Markov-chain apparatus output on parameters read from the published biochemistry or from structural homology. *Predicted* = framework-derived experimental target not yet measured.

| Claim | Status | Source / basis |
| --- | --- | --- |
| Drt3b produces alternating poly(AC) without nucleic acid template | Observed | Deng *et al.* [5] biochemistry |
| Glu26-to-Gln produces 80%/20% dA/dG split at A-state | Observed | Deng *et al.* [5, Fig. 4J] |
| AbiK same-fold produces random ssDNA | Observed | Deng *et al.* [5] |
| 1,232-homolog alignment with Glu26, Gly248, Arg253 conservation pattern | Observed | Deng *et al.* [5] Data S3 |
| Marginal G fraction $\approx 0.10$ in Glu26-to-Gln (analytical $0.5 \times 0.20$) | Derived | Half-cycle dG misincorporation arithmetic |
| Period-2 self-match $\approx 0.857$ in Glu26-to-Asp clades | Derived | $\frac{1}{2} \sum p_A^2 + \frac{1}{2} \sum p_C^2$ from parameterization |
| Mode 3 capacity ceiling $\log_2 N$ | Derived | Capacity theorem applied to $N$-state cyclic substrate |
| WT Drt3b: $I_{\text{struct}}^{\text{pop}} = 1.0$ bits, $f_{\text{phase}} \approx 0.99$, separation ratio $234 \times - 1031 \times$ | Simulated | Apparatus on Markov channel parameterized from observed (AC) output |
| Glu26-to-Gln: $I_{\text{struct}}^{\text{pop}} = 1.0$ bits, $f_{\text{monomer}} \approx 0.80$ A-state | Simulated | Apparatus on Markov channel parameterized from observed 80%/20% split |
| 33/33 secondary-residue clades in-envelope | Simulated | Apparatus on per-clade Markov parameterizations from residue assignments |
| 15.7-fold elevated dG in 6 Glu26-to-Asp clades | Simulated | Apparatus on per-clade parameterizations; not measured from raw products |
| Drt3a vs Drt3b separation $\Delta I_{\text{struct}}^{\text{chan}} = 1.0$ bit | Simulated | Channel-MI against stated 5-channel ensemble |
| Arg253-to-Ala: $I_{\text{struct}}^{\text{pop}} \rightarrow 0.009$, phase peak $\rightarrow 0.43$, cycle collapse | Predicted | SDM experimental target; not yet performed |
| Gly248-to-Ala: $I_{\text{struct}}^{\text{pop}}$ near WT, marginal G $\rightarrow 0.15$, cycle intact | Predicted | SDM experimental target; not yet performed |
| Arg253-to-Ala + Gly248-to-Ala double mutant: Arg253-to-Ala signature dominates | Predicted | SDM experimental target; test of gate-class ordering |

**TABLE 5:**
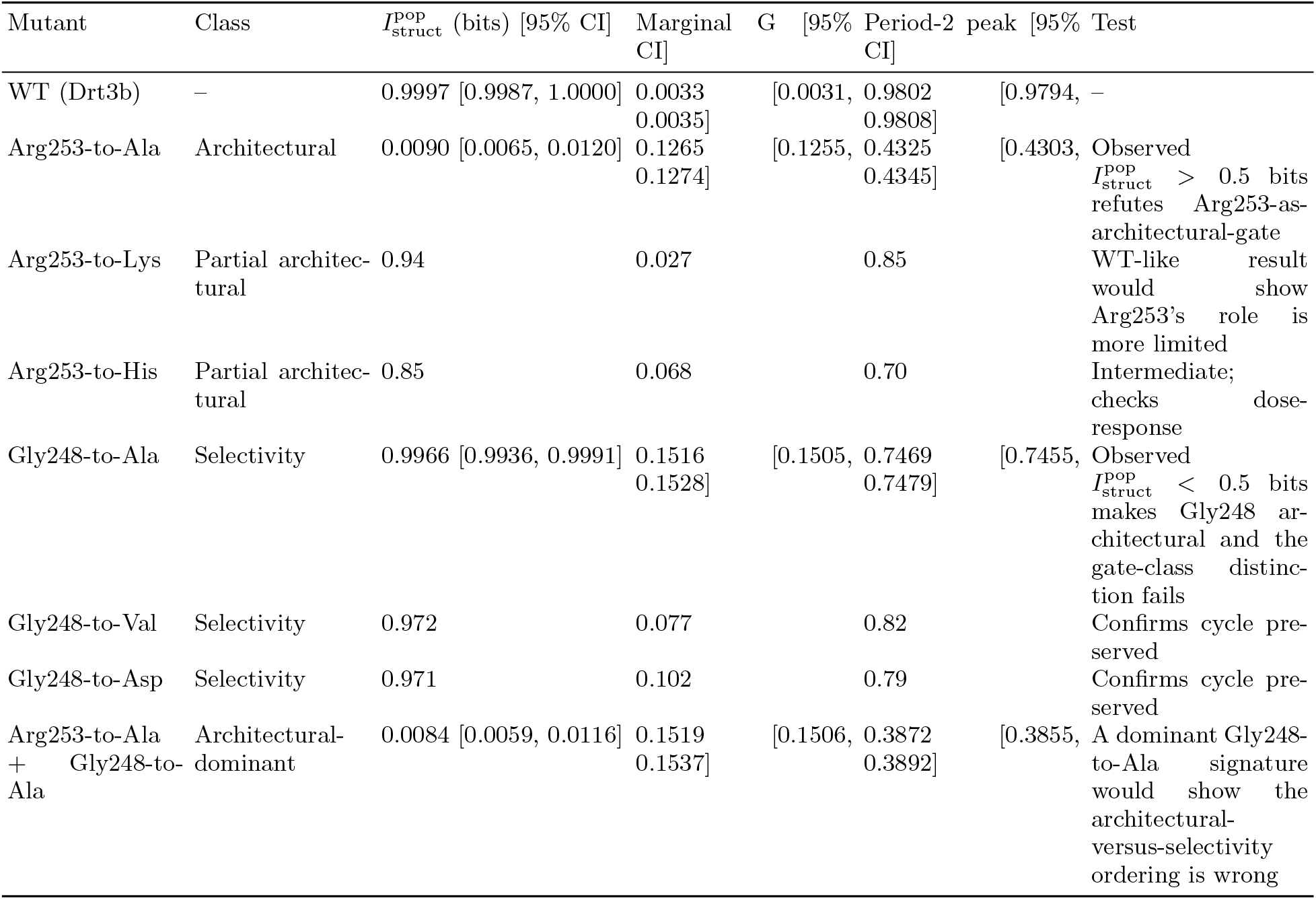
Universal-gate site-directed-mutagenesis predictions (architectural versus selectivity). Predictions at *L* = 64, 30 reps, *n*_samples_ = 5000 per rep. 95% CIs from beta-binomial.

**TABLE 6:**
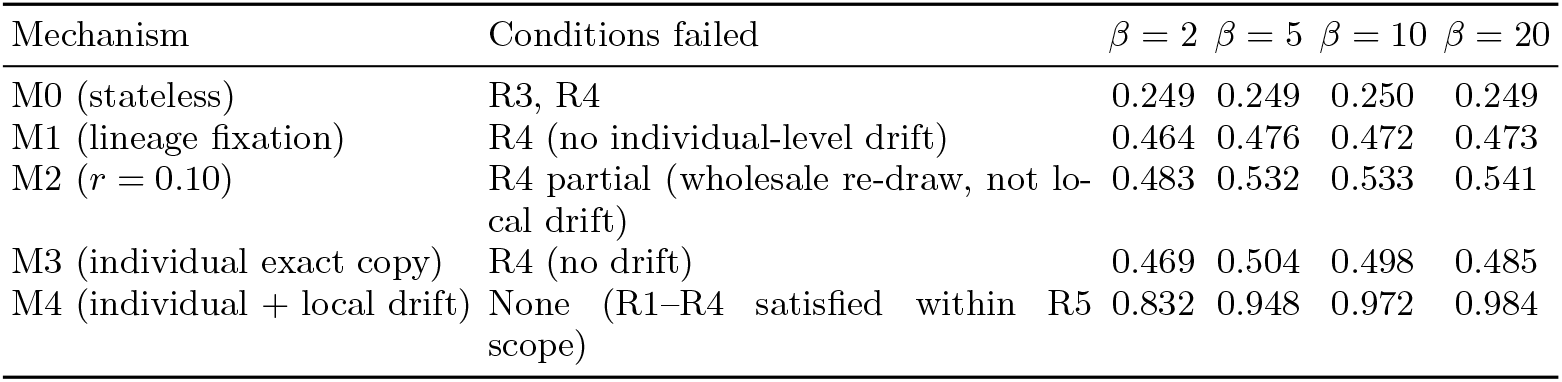
Inheritance-mechanism plateau heights (Test H3, Corollary 1). Plateau = mean fitness at gen 1000 averaged over 30 replicates. *K* = 400, *L*_TARGET_ = 32, *µ*_*M*4_ = 0.01. Theoretical maximum 1.0; chance baseline 0.250. Mechanism-condition mapping in column 2.

### H. Test F: apparatus signature on Drt3 channels

Five-channel ensemble: {Drt3a Mode 1, Drt3b Mode 3 WT, Drt3b Glu26-to-Gln, Drt3a homopolymer null, AbiK random}. Channel ensemble robustness: 10 alternative four-channel ensembles drawn from a 10-channel pool. Mode classification stability checked across all ensemble draws.

### I. Test G: Drt3b homolog family-level apparatus application

1,232 Drt3b homologs from Deng et al. supplementary alignment. Clade assignment by primary-gate residue (Glu26, Gln26, Asp26, other) and secondary-residue identity at six conserved positions. Apparatus signature predicted per clade; in-envelope assessment at the WTDrt3b reference signature.

### J. Test H: five-mechanism population-dynamics characterization

Mechanisms M0–M4 as defined in Results. Target length *L*_TARGET_ ∈ {16, 32, 64, 128}. Population size *K* = 400. Selection sharpness *β* ∈ {2, 5, 10, 20}. Mutation rate (M4) *µ* ∈ {0.001, 0.01, 0.05, 0.10}. Re-draw rate (M2) *r* 0.01, 0.05, 0.10, 0.25, 0.50. 30 replicates per cell. Plateau height equals mean fitness at gen 1000. Direct competition uses balanced start with equal initial population sizes; the head-to-head winner is the 5000-gen majority. Long-horizon stability assessed at 5000 generations.

### K. Test I: universal-gate SDM predictions

Arg253 substitutions: Arg253-to-Ala, Arg253-to-Lys, Arg253-to-His. Gly248 substitutions: Gly248-to-Ala, Gly248-to-Val, Gly248-to-Asp. Double mutant: Arg253to-Ala + Gly248-to-Ala. Markov-chain channels parameterized from structural homology arguments and the Drt3b WT reference. *L* = 64, 30 replicates, *n*_samples_ = 5000 per replicate. 95% CIs from beta-binomial.

## DATA AVAILABILITY

All simulation code, raw simulation outputs, the diagnostic apparatus implementation, the family-level analysis pipeline, and the dispatch markdown files containing the pre-specified analysis plans for each test cohort are deposited in a permanent public repository (github.com/khatvangi/templating-substrates) and archived on Zenodo with two complementary DOIs. 10.5281/zenodo.20060972 is the *pre-specification anchor* (release v0.0-prereg), archiving the analysis-plan dispatches at the moment of pre-registration before the empirical work began. 10.5281/zenodo.20272479 is the *reproducibility snapshot* (release v1.0-submission), archiving the full submission state including manuscript, supplementary appendix, bibliography, figures, all test scripts, and all result CSVs. Phylogenetic input files (`science aed1656 data s4--s8`) are derived from the published supplementary material of Deng *et al*. [5]. The deposit’s git history records the commit ordering: Phase 0 retroactively commits the dispatches with their original filesystem mtimes preserved as commit metadata, and Phases 1–5 commit the test scripts, result CSVs, and figure pipeline. The pre-specified predictions are reproduced verbatim in the docstring headers of the test scripts committed in Phases 2–4.

## Footnotes

1 Analysis plans for each test cohort were written in dispatch markdown files (`dispatch1.md` through `dispatches v4/`) and reproduced verbatim in the docstring header of each test script before the corresponding result CSVs were generated. The full deposit is archived at 10.5281/zenodo.20060972 (Phase 0 retroactively committing the dispatches; Phases 1–5 adding scripts and result CSVs).

