## Supplementary material for "A substrate recursion principle for biological information, with empirical anchoring through a templating-mode taxonomy": supplemntal

### Supplementary Mathematical Appendix
### A theory of templating substrates

Boggavarapu Kiran

This appendix provides formal definitions and proofs for the mathematical claims in the main paper. The material is organized into five sections corresponding to the apparatus, mode-specific information bounds, descriptor-relativity, the formal sense of bulk-matched controls, and the capacity and generation theorems with their bounded-heredity corollary and finite-population proposition.

The appendix is written for a reader with graduate-level information theory (Cover and Thomas 2006) and basic familiarity with discrete memoryless channels. Notation follows the main paper; $|\mathcal{A}|$ denotes alphabet cardinality, $h_b(p) = -p\log_2 p - (1-p)\log_2(1-p)$ is binary entropy, and $D_{\mathrm{KL}}(P \parallel Q)$ is Kullback-Leibler divergence.

#### Formal apparatus definitions

##### The templating event

A templating event is a tuple $\mathcal{E} = (X, O, S, \Delta G; \varphi_X, \varphi_Y, Y)$ where:

- $X \in \mathcal{X}$ is the template, drawn from a template population with distribution $\Pi_X$ (which may be a point mass $\delta_x$ for a single fixed template).
- $O$ is the operator (separable molecular machinery; possibly empty if templating is autocatalytic).
- $S$ is the substrate pool from which $Y$ is constructed.
- $\Delta G$ is the free-energy budget driving the templating reaction.
- $\varphi_X : \mathcal{X} \to \mathcal{X}'$ and $\varphi_Y : \mathcal{Y} \to \mathcal{Y}'$ are descriptor maps on template and product spaces.
- $Y$ is the structured product, distributed as $Y \mid (X, O, S, \Delta G) \sim P_{\mathrm{chan}}(\cdot \mid X)$, where $P_{\mathrm{chan}}$ is the templating channel induced by $O$ acting on $(X, S, \Delta G)$.

The templating channel $P_{\mathrm{chan}}(\cdot \mid X)$ is a stochastic kernel from $\mathcal{X}$ to $\mathcal{Y}$. When $\varphi_X = \varphi_Y$ the templating is *same-descriptor*; otherwise it is *cross-descriptor* and $O$ implements the descriptor coupling.

#### Population mutual information

**Definition 1** (Population MI). For a templating event $\mathcal{E}$ with template population $\mathcal{X}$ and distribution $\Pi_X$,

$$I_{\text{struct}}^{\text{pop}}(\mathcal{E}) := I(\varphi_X(X); \varphi_Y(Y)) = H(\varphi_Y(Y)) - H(\varphi_Y(Y) \mid \varphi_X(X)),$$

where the marginal $H(\varphi_Y(Y))$ is computed under the law $Y \sim \int_{\mathcal{X}} P_{\text{chan}}(\cdot \mid x)\, d\Pi_X(x)$ and the conditional entropy is over the joint law of $(\varphi_X(X), \varphi_Y(Y))$ induced by $\Pi_X$ and $P_{\text{chan}}$.

The definition is descriptor-relative: it measures information transfer between the *declared descriptors* $\varphi_X$ and $\varphi_Y$, not between raw $X$ and $\varphi_Y(Y)$. Conditioning on full $X$ would leak structure outside the declared descriptor and undermine the framework's descriptor-relativity claims (§3); the form above is consistent with the empirical estimator, which takes $(\varphi_X(X), \varphi_Y(Y))$-paired samples and never reads $X$ outside $\varphi_X$.

**Proposition 2** (Population MI bound). $I_{\text{struct}}^{\text{pop}}(\mathcal{E}) \le H(\varphi_X(X)) \le H(\Pi_X)$.

*Proof*. The first inequality is $I(\varphi_X(X); \varphi_Y(Y)) \le H(\varphi_X(X))$, the standard MI–entropy bound. The second is data-processing applied to $\varphi_X$ as a deterministic post-processing of $X$: $H(\varphi_X(X)) \le H(X) = H(\Pi_X)$. □

**Corollary 3** (Fixed-template degeneracy). *When $\mathcal{X} = \{x_0\}$ is a single fixed template, $H(\Pi_X) = 0$, so $I_{\text{struct}}^{\text{pop}}(\mathcal{E}) = 0$ by construction.*

This corollary is the formal content of the framework's claim that biological systems with single fixed templates (Drt3a's single ncRNA, a single chromosome being replicated within a cell) have zero population MI. Population MI measures *cross-realization information transfer*; for a fixed template, there are no realizations to transfer between.

#### Substrate capacity

The capacity of a templating substrate is defined as a supremum of population MI over input ensembles. Because the framework's heredity claim concerns *recursive* information transfer (the product must itself be readable as a next-round template), the supremum is taken over ensembles supported on the recursively closed set:

$$\mathcal{R}_L := \{X \in \mathcal{X}_L : \mathcal{O}^t(X) \in \mathcal{X}_L \text{ for all } t \ge 0, \text{possibly via the reversible involution } \iota\}.$$

Restricting the supremum to $\mathcal{R}_L$ makes R3 (catalytic closure under iteration) mandatory in the capacity definition rather than argued for separately in the proof. Substrates failing R3 (e.g., translation, where the protein product is not ribosome-readable) have $\mathcal{R}_L = \emptyset$ or trivially small, so their substrate capacity is zero or bounded by a single-application transfer.

**Definition 4** (Substrate capacity, recursive). For a templating substrate class $\mathcal{S}$ and template length $L$, the substrate capacity is

$$C_{\mathcal{S}}(L) := \sup_{\Pi_X \text{ on } \mathcal{R}_L} I_{\text{struct}}^{\text{pop}}(\mathcal{E}).$$

The supremum is over template-distribution choices on the recursively closed input set $\mathcal{R}_L \subseteq \mathcal{X}_L$. $C_{\mathcal{S}}$ is a property of the substrate class, not of any particular biological realization. In contrast, $I_{\text{struct}}^{\text{pop}}$ is the realized population MI in a specified system. Both are needed; conflating them is the most common failure mode of substrate-level reasoning.

**Remark 5** (On the choice to restrict to $\mathcal{R}_L$). A reader might prefer to define a separate one-step channel capacity over all of $\mathcal{X}_L$. That object is well-defined and useful for non-recursive transfers (e.g., translation), but it cannot ground a *recursive* heredity claim: a substrate with $\Omega(L)$ one-step capacity but $\mathcal{R}_L = \emptyset$ transmits information one time and then exits the substrate class. Heredity, in the framework's sense, requires the iteration to remain within $(\mathcal{X}, \mathcal{O})$, and this is exactly what restricting to $\mathcal{R}_L$ encodes.

**Remark 6**. For Mode 1 (Watson-Crick) at alphabet size $|\mathcal{A}|$ and per-position misincorporation rate $\varepsilon$, $C_{\mathcal{S}}(L) = L \cdot [\log_2|\mathcal{A}| - h_b(\varepsilon) - \varepsilon \log_2(|\mathcal{A}| - 1)]$, achieved by the uniform distribution $\Pi_X^{\text{unif}}$ on $\mathcal{R}_L$ (which for Mode 1 equals $\mathcal{X}_L$ up to involution-equivalence). For Mode 3 with $N$-state cyclic dynamics, $C_{\mathcal{S}}(L) \le \log_2 N$ for all $L$, regardless of template-distribution choice (Theorem 14).

#### Channel mutual information

**Definition 7** (Channel MI against an ensemble). Let $\mathcal{C} = \{c_1, \dots, c_K\}$ be a finite ensemble of templating channels with prior $P(C)$. For a focal channel $c \in \mathcal{C}$,

$$I_{\text{struct}}^{\text{chan}}(c; \mathcal{C}) := D_{\text{KL}}(P(\varphi_Y(Y) \mid C = c) \parallel \bar{P}(\varphi_Y(Y)))$$

where the mixture distribution is

$$\bar{P}(\varphi_Y(Y)) := \sum_{c' \in \mathcal{C}} P(C = c') P(\varphi_Y(Y) \mid C = c').$$

The full ensemble channel MI is

$$\bar{I}_{\text{struct}}^{\text{chan}}(\mathcal{C}) := \sum_{c \in \mathcal{C}} P(C = c) D_{\text{KL}}(P(\varphi_Y(Y) \mid C = c) \parallel \bar{P}).$$

**Proposition 8** (Channel MI bound). $\bar{I}_{\text{struct}}^{\text{chan}}(\mathcal{C}) \le \log_2|\mathcal{C}|$, *with equality iff the channels in* $\mathcal{C}$ *produce mutually disjoint output distributions.*

*Proof.* $\bar{I}_{\text{struct}}^{\text{chan}}(\mathcal{C}) = I(C; Y)$ under the joint law $P(C, Y) = P(C)P(Y \mid C)$. $H(C) \le \log_2|\mathcal{C}|$. The mutual information is bounded by either marginal entropy. Equality of $I(C; Y)$ and $H(C)$ occurs iff $Y$ determines $C$, i.e., iff the conditional distributions $P(Y \mid C = c)$ have disjoint supports. ☐

**Remark 9** (Channel MI for fixed templates). Unlike $I_{\text{struct}}^{\text{pop}}$, $I_{\text{struct}}^{\text{chan}}$ does not vanish for fixed-template channels. A channel with a single fixed template still produces a distinguishable output distribution against alternative channels, and the divergence $D_{\mathrm{KL}}(P(Y \mid C = c) \parallel \bar{P})$ measures that distinguishability. This is the formal content of the framework's claim that $I_{\mathrm{struct}}^{\mathrm{chan}}$ distinguishes Drt3a (Mode 1 with fixed ncRNA template) from Drt3b (Mode 3 with cyclic active site) even when both are individual cellular realizations with fixed mechanisms.

#### Per-base mechanism fidelity

**Definition 10** (Phase fidelity and monomer fidelity). For a templating channel with cyclic dynamics having $N$ states, define a phase-trajectory function $\sigma: \mathbb{N} \to \{0,1,\dots,N-1\}$ that gives the canonical phase for each position. The *phase fidelity* is

$$f_{\mathrm{phase}}(c) := \mathbb{E}\left[\frac{1}{L}\sum_{i=1}^{L} \mathbf{1}\{\hat{\sigma}_i(Y) = \sigma(i)\}\right]$$

where $\hat{\sigma}_i$ is the phase recovered from position $i$ of $Y$. The *monomer fidelity* at state $s$ is

$$f_{\mathrm{monomer}}(c; s) := \mathbb{E}\left[\frac{1}{|\{i: \sigma(i) = s\}|}\sum_{i:\sigma(i)=s} \mathbf{1}\{Y_i = \pi(s)\}\right]$$

where $\pi(s)$ is the canonical monomer for state $s$. For acyclic channels (Mode 1, Mode 5), $N = 1$, $f_{\mathrm{phase}} \equiv 1$, and only $f_{\mathrm{monomer}}$ is meaningful, simplifying to the per-position correctness rate.

**Remark 11**. $f_{\mathrm{phase}}$ and $f_{\mathrm{monomer}}$ are not mutual informations. They are moments of the joint distribution capturing different mechanism-level observables. The split is necessary because cyclic channels can fail in two structurally distinct ways: an architectural disruption that breaks $f_{\mathrm{phase}}$ (the cycle no longer alternates) versus a selectivity disruption that degrades $f_{\mathrm{monomer}}$ within an intact cycle (the alternation continues but with degraded monomer choice). These correspond to the architectural-versus-selectivity gate distinction reported in main paper Results.

#### The apparatus signature

**Definition 12** (Joint signature). The apparatus signature for a templating channel $c$ against an ensemble $\mathcal{C}$ is the tuple

$$\sigma(c; \mathcal{C}) := \left(I_{\mathrm{struct}}^{\mathrm{pop}}(c), I_{\mathrm{struct}}^{\mathrm{chan}}(c; \mathcal{C}), f_{\mathrm{phase}}(c), f_{\mathrm{monomer}}(c; s)\right)_{s=0}^{N-1}.$$

Mode classification uses the joint signature; signature collapse to any single coordinate is information-discarding.

### Mode-specific information bounds

This section proves the scaling-law claims for each of the five primary modes. Mode 6 is treated separately in §2.6.

#### Mode 1 — linear scaling

**Theorem 13** (Mode 1 capacity). *For a Watson-Crick channel with alphabet $|\mathcal{A}| \geq 2$, per-position misincorporation rate $\varepsilon$, template population $\mathcal{X} = \mathcal{A}^L$ with uniform distribution $\Pi_X^{\mathrm{unif}}$, and product $Y$ of length $L$, $I_{\mathrm{struct}}^{\mathrm{pop}}(X;Y) = L \cdot [\log_2|\mathcal{A}| - h_b(\varepsilon) - \varepsilon \log_2(|\mathcal{A}| - 1)]$.*

*Proof.* The Watson-Crick channel is discrete memoryless: $P(Y_i = \mathrm{wc}(x_i) \mid X_i = x_i) = 1 - \varepsilon$, and $P(Y_i = a' \mid X_i = x_i) = \varepsilon/(|\mathcal{A}| - 1)$ for each $a' \neq \mathrm{wc}(x_i)$. By memorylessness, the channel decomposes per position. For any position $i$, $I(X_i;Y_i) = H(X_i) - H(X_i \mid Y_i)$. Under uniform $\Pi_X$, $H(X_i) = \log_2|\mathcal{A}|$. By Bayes' rule and uniformity, $H(X_i \mid Y_i) = H(Y_i \mid X_i) = h_b(\varepsilon) + \varepsilon \log_2(|\mathcal{A}| - 1)$. Summing over $L$ positions gives the stated formula. □

*Empirical match.*

Across 30 sweep cells with $\varepsilon \in \{0.001, 0.01, 0.05, 0.10, 0.25\}$ and $L \in \{10, 25, 50, 100, 200, 500\}$, $n_{\mathrm{samples}} = 5000$ per cell, the empirical $I_{\mathrm{struct}}^{\mathrm{pop}}$ matches Theorem 13 within 0.6% across all cells (Test A.1, main paper Fig. 1A).

#### Mode 3 — logarithmic saturation

**Theorem 14** (Mode 3 capacity bound). *For an $N$-state cyclic templating channel with phase $X \in \{0, 1, \dots, N-1\}$ drawn uniformly, and product $Y$ of length $L$ generated by deterministic cycle progression with per-step substrate-selection error $\varepsilon$, $I_{\mathrm{struct}}^{\mathrm{pop}}(X;Y) \leq \log_2 N$ for all $L$, provided the $N$ phase states are pairwise distinguishable under the product descriptor $\varphi_Y$ (so that the phase-conditioned output distributions $P(\varphi_Y(Y) \mid X = j)$ are pairwise distinct). The bound is approached in the low-noise long-block limit ($L \to \infty$ with $\varepsilon$ fixed at small positive value); the bound is saturated exactly at $\varepsilon = 0$.*

*Proof.* By the data-processing inequality applied to the cycle progression (which is a deterministic function of phase and time step), $I(X;Y) \leq I(X; \text{phase trajectory}) \leq H(X) = \log_2 N$. The bound is achieved iff phase is recoverable from $Y$, which requires the phase-conditioned output distributions to be pairwise distinguishable. If two phase states $j \neq j'$ produce identical output distributions under $\varphi_Y$, then $I(X;Y) < \log_2 N$ strictly. Under pairwise distinguishability and $\varepsilon = 0$, phase is recoverable from $Y_1$ alone (the first position determines the phase that produced it). At $\varepsilon > 0$ the recoverability is imperfect on a single position but improves with $L$ because the periodic structure of $Y$ provides redundant phase information. Standard channel-coding asymptotic analysis (Cover and Thomas 2006, Theorem 8.7.1) gives $\lim_{L\to\infty} I(X;Y) = \log_2 N$ for fixed $\varepsilon > 0$ in the regime where the cycle's per-step Shannon capacity exceeds zero. □

*Empirical match.*

For $N \in \{2,3,4,5,6,8,10\}$, $\varepsilon = 10^{-3}$, $L$ from $L = N$ to $L = 50N$: empirical $I_{\text{struct}}^{\text{pop}}$ approaches $\log_2 N$ to within estimator precision (drift across $L$ values $< 0.0001\%$; Test B, main paper Fig. 2A,B).

#### Mode 5 — module-bounded

**Theorem 15** (Mode 5 capacity). *For a modular conveyor with $N$ modules, each independently selecting one monomer from substrate alphabet of size $k$ with fidelity $1 - \varepsilon$, and product $Y$ of length $L$: $I_{\text{struct}}^{\text{pop}}(X;Y) \leq \min(L,N) \cdot [\log_2 k - h_b(\varepsilon) - \varepsilon \log_2(k-1)]$, with positions beyond $\min(L,N)$ contributing zero information (templated positions saturate at $N$; untemplated positions in a $L > N$ product carry no per-position information about the modular template).*

*Proof.* Each module independently determines one product position; modules 1 through $\min(L,N)$ are templated, positions $N+1,\ldots,L$ (when $L > N$) have no module association and are drawn from a fixed distribution independent of the modular template. Per-position MI follows the same memoryless argument as Theorem 13 for templated positions and equals 0 for non-templated positions. Summing over the $\min(L,N)$ templated positions gives the stated bound. □

*Empirical match.*

Test C, main paper Fig. 2D.

#### Mode 2 — code-mediated capacity ceiling

**Theorem 16** (Mode 2 capacity bound). *For code-mediated translation with DNA template of $L$ codons drawn uniformly random from the 64 codons, standard genetic code (NCBI table 1), translation error rate $\varepsilon \ll 1$, and peptide output $Y$ of length $L$ over 20 amino acids plus stop, $I_{\text{struct}}^{\text{pop}}(X;Y) \leq L \cdot H(\nu)$, where $\nu$ is the codon-count distribution: $\nu(a) = n(a)/64$, with $n(a)$ the number of codons mapping to amino acid $a$ in the standard genetic code (e.g., $n(\text{Leu}) = 6$, $n(\text{Met}) = 1$).*

*Proof.* $I(X;Y) \leq H(Y)$. For uniform random $X$, the marginal distribution of $Y_i$ (per codon) is exactly $\nu$, so $H(Y_i) = H(\nu)$. By memorylessness of per-codon translation, $H(Y) = L \cdot H(\nu)$. □

*Empirical match.*

$H(\nu) = 4.218$ bits/codon for the standard genetic code (computed from the codon table, including stop as a 21st symbol); empirical $I_{\text{struct}}^{\text{pop}}$ at $L = 64$, $n_{\text{samples}} = 1000$, gives 4.214 bits/codon (Test F4). Match to four significant figures.

*Capacity comparison with Mode 1.*

The same DNA at length $3L$ nucleotides has Mode 1 capacity $6L$ bits (Theorem 13 at $|\mathcal{A}| = 4$). Mode 2 capacity is $L \cdot H(\nu) = 4.218L$ bits. Capacity ratio $= 6/4.218 \approx 1.42$. Code degeneracy reduces information transfer by approximately 30% from the substrate ceiling.

*Note on alternative bounds.*

The frequently-cited $L \cdot \log_2(20) = 4.32L$ bits assumes uniform marginal over amino acids, which the standard genetic code does not produce; the correct bound uses $H(\nu)$. Including stop as a 21st symbol gives $H(\nu) = 4.218$ bits; restricting to the 61 sense codons only gives $H(\nu_{\text{sense}}) \approx 4.139$ bits.

*Low-error caveat.*

The bound $L \cdot H(\nu)$ is the no-error / low-error ceiling: it assumes translation fidelity is high enough that codon-to-amino-acid assignment is effectively deterministic. With non-negligible mistranslation rate $\varepsilon_t > 0$ per codon, the per-codon channel introduces a noisy mapping with capacity strictly below $H(\nu)$, and the realized $I_{\text{struct}}^{\text{pop}}$ is correspondingly lower. The theorem above states the ceiling; including mistranslation requires a per-codon channel-capacity correction analogous to the $h_b(\varepsilon)$ term in Theorem 13.

#### Mode 4 — length independence

**Theorem 17** (Mode 4 capacity bound). *For a conformer-state templating channel with $K$ distinguishable conformers and templated polymers of length $L$, $I_{\text{struct}}^{\text{pop}}(X;Y) \leq \log_2 K$ independently of $L$.*

*Proof.* $X$ (the templating conformer) takes $K$ values; $H(X) \leq \log_2 K$. The product $Y$ inherits the conformer assignment as a single discrete attribute (its conformational state), regardless of polymer length. The polymer's primary structure is fixed by the genome, not by the conformer. □

**Remark 18**. Mode 4 is “state-templating” rather than “position-templating”; the length-independence is a property of the substrate (conformational identity), not a degenerate special case of position templating. The framework treats Mode 4 as $O(1)$ information content as a structural feature, not as an asymptotic limit.

#### Mode 6 — surface-position capacity is possible; biological instances are parasitic and R4-limited

For a 2D surface-position templating system with $N$ distinguishable surface positions and per-position selection from $k$ options, the combinatorial capacity bound is

$$C_S^{\text{combinatorial}}(N) \leq N \cdot \log_2 k,$$

which can in principle scale unboundedly as the surface area grows. Mode 6 capacity scaling is therefore not bounded by dimensionality alone.

The framework's distinguishing claim about Mode 6 is that biological instances fail two operational requirements distinct from raw combinatorial capacity:

1. **Parasitic copyability.** The operator that copies the surface pattern, and the units that the surface assembles, are both produced by Mode 1 acting on the genome. Failure of Mode 1 inheritance terminates Mode 6 inheritance. Mode 6's apparent capacity is gated by Mode 1's substrate.

2. **Bounded heritable variation.** The variation that biological Mode 6 transmits is geometric perturbation of an existing unit repertoire, not compositional novelty. The number of distinguishable cortical states a lineage can transmit is bounded by combinatorial arrangements of genome-encoded parts, not by the surface's combinatorial position count. Open-ended scaling requires unboundedly many heritable distinguishable states; biological Mode 6 lacks this because new compositional novelty would require new genome-encoded parts.

We omit a formal bound because Mode 6's biological instances (ciliate cortex) are not fully characterized at the apparatus level in the published literature. The operational implementation of bounded heredity in the H3 mechanism characterization is the empirically relevant claim; see §5 below.

### Descriptor-relativity

**Definition 19**. Two descriptor maps $\varphi_Y, \varphi'_Y$ on $\mathcal{Y}$ are *equivalent* if there exists a bijection $\pi$ such that $\varphi'_Y = \pi \circ \varphi_Y$. A templating event $\mathcal{E}$ analyzed under non-equivalent descriptors $\varphi_Y$ and $\varphi'_Y$ may give different $I_{\text{struct}}^{\text{pop}}$ values.

**Theorem 20** (Descriptor non-uniqueness for designed-replicator templating). *Let a templating event $\mathcal{E}$ propagate constraints between a parent and daughter unit through residue-pairing chemistry (e.g., a designed $\beta$-sheet peptide replicator in which residues at paired positions select their partners by side-chain complementarity). Under a descriptor $\varphi_Y^{\text{pair}}$ that records residue identity at each paired position, the templating event classifies as Mode-1-like with information bound determined by the pairing constraints, and $I_{\text{struct}}^{\text{pop}}$ scales with the number of paired positions. Under a descriptor $\varphi_Y^{\text{conf}}$ that records only the conformational state (which $\beta$-sheet topology was adopted), the same templating event has $I_{\text{struct}}^{\text{pop}} = O(1)$.*

*Sketch*. Under $\varphi_Y^{\text{pair}}$, the joint distribution of daughter-unit residue identities given parent residue identities at paired positions has the structure of a position-by-position channel with per-position capacity determined by the pairing-rule alphabet; $I_{\text{struct}}^{\text{pop}}$ scales linearly with the number of paired positions, subject to those constraints. Under $\varphi_Y^{\text{conf}}$, only the conformational state is observed; the daughter inherits the parent's conformation regardless of how many paired positions exist, so $I_{\mathrm{struct}}^{\mathrm{pop}}$ is bounded by $\log_2(\text{number of distinguishable conformations}) = O(1)$. □

**Remark 21** (Why not amyloid sequence-templating). A naive version of this point invokes amyloid or prion sequence propagation as the classifier-dependent case. That framing is incorrect: in biological prions, the amino-acid sequence is genome-encoded, and the amyloid does not template primary sequence at all. The phenomenon biological amyloids transmit is conformational — not sequence — and the framework treats this as a clean Mode 4 case (see Discussion, yeast-prion paragraph). The descriptor-relativity point made here is a property of templating systems where pairing-chemistry constraints *do* act position-by-position on the daughter unit's primary structure; designed $\beta$-sheet replicators and analogous synthetic systems are the cleanest examples.

**Corollary 22**. *Mode classification applies to (templating event, descriptor) pairs, not to physical objects. The same template can mediate templating events that classify into different modes under different descriptors.*

This is what allows the framework to avoid an unresolvable taxonomic question of the form "is amyloid templating Mode 1 or Mode 4?" The question is ill-posed; the answer depends on what is being measured.

### Bulk-matched controls as null hypotheses

**Definition 23** ($k$-th order matched null). Given a product $Y$ from a templating event, the $k$-th order matched null distribution $Q_k$ is the maximum-entropy distribution on $\mathcal{Y}$ that preserves all $k$-th order marginal statistics of the empirical distribution of $Y$. Specifically: $Q_1$ preserves single-position marginals (composition) and is iid from those marginals; $Q_2$ preserves nearest-neighbor frequencies and is the maximum-entropy first-order Markov chain consistent with those frequencies; $Q_k$ preserves $k$-position joint marginals and is the maximum-entropy $(k-1)$-th order Markov chain.

**Remark 24**. For $k = 2$, $Q_2$ is *not* iid; samples from $Q_2$ are first-order Markov sequences whose nearest-neighbor frequencies match the empirical $Y$.

**Definition 25** (Bulk-matched paired null). The bulk-matched paired null for a templating event $\mathcal{E}$ is constructed by holding the template ensemble fixed and replacing the channel output with a draw from a $k$-th order matched distribution that destroys position-specific template–product correspondence. Specifically, for each observed template $X$, the null product $Y_{\mathrm{bulk}}$ is drawn from

$$Y_{\mathrm{bulk}} \sim Q_k(\cdot \mid \text{compositional class of } X)$$

where $Q_k$ is the $k$-th order matched distribution fitted to the product ensemble, optionally conditioned on the low-order composition class associated with $X$ but *not* on $X$'s position-specific structure. This preserves $k$-th order marginal statistics between $X$ and $Y_{\mathrm{bulk}}$ while destroying the templating correspondence.

**Definition 26** (Structure-scrambled null). The structure-scrambled null is a second-order companion to Definition 25: for each $(X, Y)$ pair, replace $Y$ with $Y_{\text{scr}}$, a positionally permuted copy of $Y$ that preserves the compositional histogram but randomizes the position assignment. Position-specific information transfer from $X$ to $Y$ is destroyed; compositional information is preserved.

**Proposition 27** (Bulk and structure-scrambled controls as nulls). *A claim that $\mathcal{E}$ is a templating event with position-specific structural information transfer requires $I^{\text{pop}}_{\text{struct}}(\varphi_X(X); \varphi_Y(Y)) > \tau \cdot I^{\text{pop}}_{\text{struct}}(\varphi_X(X); \varphi_Y(Y_{\text{bulk}}))$ and $I^{\text{pop}}_{\text{struct}}(\varphi_X(X); \varphi_Y(Y)) > \tau \cdot I^{\text{pop}}_{\text{struct}}(\varphi_X(X); \varphi_Y(Y_{\text{scr}}))$ where $\tau$ is a stated significance threshold (the main paper uses $\tau \geq 20$ for a "pass" verdict).*

The threshold $\tau$ encodes the strength of the claim. The paired-null formulation is essential: in an earlier formulation, $X_{\text{bulk}}$ and $Y_{\text{bulk}}$ were drawn independently, which makes $I = 0$ exactly (modulo finite-sample estimator bias), giving a useless control. The paired null keeps the template marginal but breaks the position-specific correspondence, so any residual mutual information reflects compositional or low-order structure shared between $X$ and the product ensemble. Whether a $k$-th order match is sufficient depends on what $k$ captures the confounders: for Mode 1 versus random output, $k = 1$ is sufficient; for Mode 3 versus Mode 1 with periodic-like template, $k \geq 2$ is needed because nearest-neighbor structure differs. The framework reports controls with $k = 2$ by default.

### Capacity and generation theorems

This section formalizes the two necessity results referenced in the main paper: the *capacity theorem* (R1, R2, R3 are necessary for unlimited heredity capacity) and the *generation theorem* (R4 is additionally necessary for cumulative generative recursion beyond deterministic closure). The structure is: notation and structural conditions (§5.1); formal definition of heredity capacity with its upper bound (§5.2); formal definition of deterministic closure and cumulative generative recursion (§5.3); the capacity theorem with proof (§5.4); the generation theorem with proof (§5.5); the bounded-heredity corollary recovering each mode's capacity ceiling (§5.6); the finite-population, finite-horizon proposition (§5.7); and the recovery of prior formulations within this structure (§5.8).

#### Notation and structural conditions

Let $\mathcal{X}_L$ denote the space of admissible configurations at size parameter $L$; this is countable in all biological examples we treat. Let $\Sigma$ be a finite alphabet, called the *recognition alphabet* of the catalytic operation. A descriptor is a map $\varphi_X: \mathcal{X}_L \to \Sigma^{N(L)}$ for some position count $N(L) \geq 1$. The catalytic operation $\mathcal{O}$ has kernel $Q_L: \mathcal{X}_L \times \mathcal{X}_L \to [0,1]$ giving the conditional probability of producing $X' = \mathcal{O}(X)$ from template $X$:

$$Q_L(X' \mid X) \geq 0, \qquad \sum_{X' \in \mathcal{X}_L} Q_L(X' \mid X) = 1 \text{ for all } X.$$

We allow $\mathcal{O}$ to be composed with a reversible involution $\iota: \mathcal{X}_L \to \mathcal{X}_L$ (e.g., Watson–Crick complementation), with the recursion taking the form $X \to \iota(X') \to X'' \to \cdots$; for notational simplicity we suppress $\iota$ where it does not affect the argument.

The five conditions of the substrate recursion framework are:

**(R1) Recognition-alphabet entropy.**
$|\Sigma| \geq 2$, so $\log_2|\Sigma| > 0$.

**(R2) Unbounded extensible recognition.**
$\liminf_{L\to\infty} N(L) = \infty$, where $N(L)$ is the count of independently variable recognized positions of the descriptor $\varphi_X$ under $Q_L$. "Independently variable" is meant in the first-order, descriptor-relative sense: the substrate is not constrained to a finite recurrent phase repertoire, and increasing $L$ increases the count of heritable recognized degrees of freedom. The condition does not require strong joint-statistical independence across positions; context-dependent error rates (as in real DNA polymerases) are permitted.

**(R3) Catalytic closure under iteration.**
For every $X \in \mathcal{X}_L$ in the support of $Q_L$, $\mathcal{O}(X) \in \mathcal{X}_L$ (possibly via the reversible involution $\iota$), so that $\mathcal{O}^2(X)$ is well-defined and the iteration $X \to \mathcal{O}(X) \to \mathcal{O}^2(X) \to \cdots$ remains within $\mathcal{X}_L$.

**(R4) Stochastic drift in $\Sigma$.**
The kernel $Q_L(\cdot \mid X)$ has $|\mathrm{supp}\, Q_L(\cdot \mid X)| \geq 2$ for at least some $X \in \mathcal{X}_L$, with the dispersion of $\varphi_X(X')$ around $\varphi_X(X)$ lying in the recognition alphabet $\Sigma$ of (R1). Drift in a separate alphabet $\Sigma' \neq \Sigma$ (e.g., chemical modifications outside the polymerase's recognition palette) fails R4.

**(R5) Scope: intrinsic catalysis.**
The kernel $Q_L$ is determined by intrinsic physicochemistry — the active-site free-energy landscape, the available reactants, the chemical driving — not by an externally specified loss function or target product. R5 is scope-defining rather than structural: it demarcates the domain in which R1–R4 apply rather than introducing a fifth necessity condition.

#### Heredity capacity

Let $\Pi_X$ denote any probability distribution on $\mathcal{X}_L$ (an input ensemble of templates), and let $X' \sim Q_L(\cdot \mid X)$ for $X \sim \Pi_X$.

**Definition 28** (Heredity capacity). The *heredity capacity* of $(\mathcal{X}, \mathcal{O})$ at size $L$ is

$$C_S(L) = \sup_{\Pi_X} I(\varphi_X(X);\ \varphi_X(X')),$$

the supremum of mutual information between input and output descriptors over all input ensembles.

**Proposition 29** (Capacity upper bound). *For any kernel $Q_L$, $C_S(L) \leq N(L) \cdot \log_2|\Sigma|$.*

*Proof.* By the standard identity $I(A;B) = H(B) - H(B \mid A) \le H(B)$, for any input ensemble $\Pi_X$,

$$I(\varphi_X(X); \varphi_X(X')) \le H(\varphi_X(X')).$$

The descriptor $\varphi_X(X')$ takes values in $\Sigma^{N(L)}$, a set of cardinality $|\Sigma|^{N(L)}$, so

$$H(\varphi_X(X')) \le \log_2|\Sigma|^{N(L)} = N(L)\log_2|\Sigma|.$$

The supremum over $\Pi_X$ inherits the bound. □

The bound is achievable iff some $\Pi_X$ realizes a uniform $\varphi_X(X')$ on $\Sigma^{N(L)}$. Realizability requires that the recognition step actually resolve all $|\Sigma|$ states at each of $N(L)$ positions and that the iteration close within $\mathcal{X}_L$ — i.e., R1, R2, R3.

**Definition 30** (Unlimited heredity capacity). $(\mathcal{X}, \mathcal{O})$ has *unlimited heredity capacity* if $C_{\mathcal{S}}(L) = \Omega(L)$ as $L \to \infty$.

**Remark 31** (Information-theoretic and combinatorial formulations). Definition 30 is the information-theoretic statement. Its combinatorial equivalent: in the discrete noiseless limit where the upper bound of Proposition 29 is saturated, the recursively readable configuration set $\mathcal{R}_L = \{X \in \mathcal{X}_L : \mathcal{O}^t(X) \in \mathcal{X}_L \text{ for all } t \ge 0\}$ satisfies $|\mathcal{R}_L| \ge 2^{cL}$ for some $c > 0$. The two formulations are coordinated through entropy: linear scaling of $C_{\mathcal{S}}(L)$ in bits is equivalent to exponential scaling of the recursively readable state count. They describe the same fact at two levels.

#### Zero-drift closure and cumulative generation

A correct formal account of "what does R4 add beyond R1–R3" requires distinguishing the reachable set produced by the kernel's stochastic dynamics from the reachable set produced by a single canonical deterministic skeleton. We previously formulated this distinction using a union over all deterministic restrictions of $Q_L$; that formulation fails because, when $Q_L$ is stochastic, the union absorbs every positive-probability branch and the resulting "closure" equals the full positive-probability reachable set, making strict containment logically impossible. The corrected formulation uses the canonical zero-drift map $g_L$, which is already implicit in the framework's notion of a reference product $X_{\mathrm{ref}}$.

**Definition 32** (Zero-drift reference map). The *zero-drift reference map* of the catalytic operation $\mathcal{O}$ is the deterministic function $g_L : \mathcal{X}_L \to \mathcal{X}_L$ defined by

$$g_L(X) := X_{\mathrm{ref}}(X),$$

where $X_{\mathrm{ref}}(X)$ is the canonical product of $\mathcal{O}$ applied to template $X$ under zero drift (the most-probable successor in the noise-free or low-noise limit; e.g., the Watson–Crick complement for a DNA polymerase, the exact alternating product for Drt3b). For all biological substrates we treat, $g_L$ is well-defined; ties (if any) are broken canonically by the chemistry's reference output.

**Definition 33** (Zero-drift closure). The *zero-drift closure* of a set $S_0 \subset \mathcal{X}_L$ under $\mathcal{O}$ is

$$\mathrm{cl}_0(S_0) \;=\; \bigcup_{t\geq 0} g_L^t(S_0).$$

Equivalently, $\mathrm{cl}_0(S_0)$ is the orbit closure of $S_0$ under the deterministic zero-drift dynamics: the set of configurations reachable from $S_0$ if $\mathcal{O}$ produced its canonical reference output at every step with no error.

**Definition 34** (Cumulative generative recursion). $(\mathcal{X}, \mathcal{O})$ supports *cumulative generative recursion* if, for every $L$ sufficiently large and for every finite proper subset $S_0 \subsetneq \mathcal{X}_L$ with $|S_0| \ll |\mathcal{R}_L|$, the recursively readable descendant set

$$\mathcal{R}_{Q_L}^{\mathrm{rec}}(S_0) \;=\; \{X \in \mathcal{X}_L \colon \exists\, X_0 \in S_0,\, t \geq 0 \text{ with } Q_L^t(X \mid X_0) > 0 \text{ and } \mathcal{O}^s(X) \in \mathcal{X}_L \text{ for all } s \geq 0\}$$

strictly contains $\mathrm{cl}_0(S_0)$, with the additional configurations differing from elements of $\mathrm{cl}_0(S_0)$ in local positions of the descriptor $\varphi_X$ (i.e., in the alphabet $\Sigma$).

The zero-drift formulation does the work intended for the deterministic-closure definition without absorbing all stochastic branches. If $Q_L$ is itself deterministic and equal to $g_L$ (no drift), then $\mathcal{R}_{Q_L}^{\mathrm{rec}}(S_0) = \mathrm{cl}_0(S_0)$, not strict containment — correctly characterizing R4-failure. If $Q_L$ is stochastic with $\mathrm{supp}\, Q_L(\cdot \mid X) \supsetneq \{g_L(X)\}$ for some $X$ (i.e., R4 holds), then the descendant set contains off-orbit configurations not in $\mathrm{cl}_0(S_0)$, and strict containment holds.

**Remark 35** (Why this formulation is correct where the prior was not). The earlier formulation, $\mathrm{cl}_{\det}(S_0) = \bigcup_{f \in \mathcal{D}(Q_L)} \bigcup_t f^t(S_0)$ where $\mathcal{D}(Q_L)$ ranges over all deterministic restrictions of $Q_L$, fails because for any $X' \in \mathrm{supp}\, Q_L(\cdot \mid X)$ there exists $f \in \mathcal{D}(Q_L)$ with $f(X) = X'$. Hence $\bigcup_{f \in \mathcal{D}(Q_L)} \{f(X)\} = \mathrm{supp}\, Q_L(\cdot \mid X)$, and iterating gives $\mathrm{cl}_{\det}(S_0) =$ the full positive-probability reachable set. Strict containment $\mathcal{R}_{Q_L}^{\mathrm{rec}}(S_0) \supsetneq \mathrm{cl}_{\det}(S_0)$ is then impossible because $\mathcal{R}_{Q_L}^{\mathrm{rec}}(S_0) \subseteq \mathrm{cl}_{\det}(S_0)$ by construction. The zero-drift formulation picks a single canonical deterministic skeleton ($g_L$) rather than a union over all possible skeletons, restoring the desired strict-containment characterization of R4.

The counter-chain counterexample is still correctly excluded. A deterministic counter chain $\mathcal{X}_L = \{0,1\}^L$, $\mathcal{O}(X) = X + 1 \bmod 2^L$, has $Q_L = g_L$ (deterministic kernel coincides with its own zero-drift map), so $\mathcal{R}_{Q_L}^{\mathrm{rec}}(S_0) = \mathrm{cl}_0(S_0)$ for any $S_0$. The counter chain leaves $S_0$ but does not exceed $\mathrm{cl}_0(S_0)$; cumulative generation fails. The mistake the earlier "leaves $S_0$" formulation made — treating non-trivial deterministic dynamics as generative — is correctly avoided.

#### Capacity theorem

**Theorem 36** (Capacity necessity). *Within the (R5) scope, if $(\mathcal{X}, \mathcal{O})$ has unlimited heredity capacity in the sense of Definition 30, then R1, R2, and R3 hold.*

*Proof.* We prove the contrapositive: failure of any of R1, R2, R3 forces $C_S(L)$ to scale strictly slower than $\Omega(L)$.

*Case R1 fails*. If $|\Sigma| = 1$, then $\log_2|\Sigma| = 0$. By Proposition 29, $C_{\mathcal{S}}(L) \le N(L) \cdot 0 = 0$, so $C_{\mathcal{S}}(L) \equiv 0$, not $\Omega(L)$.

*Case R2 fails*. If $\liminf_{L\to\infty} N(L) = N_{\max} < \infty$, then $N(L) \le N_{\max}$ for all $L$ sufficiently large. By Proposition 29, $C_{\mathcal{S}}(L) \le N_{\max}\log_2|\Sigma|$, a constant in $L$. So $C_{\mathcal{S}}(L) = O(1)$, not $\Omega(L)$.

*Case R3 fails*. If $\mathcal{O}(X) \notin \mathcal{X}_L$ for some $X$ in the support of $Q_L$ (and not recoverable via reversible involution $\iota$), then for any input $X$ encountered along the iteration, there exists $t \ge 1$ with $\mathcal{O}^t(X) \notin \mathcal{X}_L$, so $X \notin \mathcal{R}_L$. The recursively closed set $\mathcal{R}_L$ is therefore empty (or restricted to a trivially small set of cycles, depending on the specific failure mode). By Definition 4, $C_{\mathcal{S}}(L) = \sup_{\Pi_X \text{ on } \mathcal{R}_L} I_{\text{struct}}^{\text{pop}}$ is taken over an empty or trivially small support set; the supremum is zero or bounded by a constant in $L$. The substrate's recursive heredity capacity, properly defined to exclude one-step transfers to non-recursive products, is therefore not $\Omega(L)$.

By contrapositive, $C_{\mathcal{S}}(L) = \Omega(L)$ requires R1, R2, and R3. □

**Remark 37** (R3 and one-step capacity). Theorem 36 is correct because the capacity is taken over $\Pi_X$ supported on $\mathcal{R}_L$, not all of $\mathcal{X}_L$. A reader familiar with channel-coding might object that one-step transfer can have $\Omega(L)$ mutual information even when R3 fails (translation: DNA → protein, $\Omega(L)$ bits in a single round). This is correct as a statement about one-step transfer but not about recursive heredity. The framework's claim concerns the substrate's capacity to host *iterated* information transfer within its own substrate class, which is exactly the restriction encoded in $\mathcal{R}_L$.

**Remark 38** (R4 is not required for capacity). Theorem 36 does not invoke R4. A perfect copier — $Q_L(X' \mid X) = \delta_{X',X}$, no drift — trivially satisfies $C_{\mathcal{S}}(L) = N(L)\log_2|\Sigma|$: the channel is the identity on $\mathcal{X}_L$, so $I(\varphi_X(X); \varphi_X(X')) = H(\varphi_X(X))$, and choosing $\Pi_X$ uniform on $\mathcal{X}_L$ achieves $H(\varphi_X(X)) = N(L)\log_2|\Sigma|$. This is the structural reason a hypothetical drift-free DNA polymerase would still be a high-capacity inheritance carrier — it would not innovate from a finite seed, but it would store and transmit the full $4^L$-fold configurational repertoire faithfully.

#### Generation theorem

**Theorem 39** (Generation necessity). *Within the (R5) scope, if $(\mathcal{X}, \mathcal{O})$ supports cumulative generative recursion in the sense of Definition 34, then R4 holds, in addition to the R1, R2, R3 inherited from coherence of the catalytic process.*

*Proof.* R1, R2, R3 are preconditions for cumulative generation as defined: cumulative generation requires a recognition alphabet (R1), at least one extensible position (R2 in the recognition-relevant sense), and a recursion that closes within $\mathcal{X}_L$ (R3). We focus on R4.

Suppose R4 fails. There are two cases.

*Case (a): the kernel is deterministic and coincides with $g_L$*. If $|\text{supp}\, Q_L(\cdot \mid X)| = 1$ for all $X \in \mathcal{X}_L$, then the unique positive-probability successor of $X$ is $X_{\text{ref}}(X) = g_L(X)$ (by the definition of $X_{\text{ref}}$ as the canonical zero-drift product). So $Q_L$ on its support equals $g_L$, and every positive-probability trajectory of $Q_L$ from $S_0$ is contained in the zero-drift orbit closure:

$$\mathcal{R}^{\text{rec}}_{Q_L}(S_0) \;=\; \bigcup_{t\geq 0} g_L^t(S_0) \;=\; \text{cl}_0(S_0).$$

This is not strict containment. Cumulative generation in the sense of Definition 34 fails.

*Case (b): the kernel drifts in an alphabet* $\Sigma' \neq \Sigma$. Suppose $Q_L(X' \mid X) > 0$ for some $X' \neq g_L(X)$, but the difference $\varphi_X(X') \neq \varphi_X(g_L(X))$ at position $i$ involves an alphabet character $\sigma' \in \Sigma' \backslash \Sigma$. Then $\varphi_X(X')$ is not in $\Sigma^{N(L)}$ under the recognition map $\varphi_X$: the drift is invisible to the next iteration's recognition step. The next $\mathcal{O}$-application reads $\varphi_X(X')$ as if it lay in $\Sigma^{N(L)}$, treating $\sigma'$ as the closest $\Sigma$-character or as undefined; either way, the drift in $\Sigma'$ does not propagate as recursively readable variation. Hence $X'$ with $\Sigma'$-drift is excluded from $\mathcal{R}^{\text{rec}}_{Q_L}(S_0)$ (the recursive-readability requirement). The descendant set therefore does not strictly contain $\text{cl}_0(S_0)$ *through branching local alternatives in* $\Sigma$ (Definition 34's qualifier).

In both cases, the cumulative-generation requirement fails. By contradiction, R4 holds whenever $(\mathcal{X}, \mathcal{O})$ supports cumulative generation. □

**Remark 40** (Counter-chain counterexample under the corrected definition). A deterministic counter chain illustrates why "leaves $S_0$" would have been insufficient and shows that the zero-drift formulation correctly excludes the counterexample. Take $\mathcal{X}_L = \{0,1\}^L$, $S_0 = \{0^L\}$, and $\mathcal{O}(X) = X + 1 \mod 2^L$ with $g_L = \mathcal{O}$ (the kernel is deterministic and its canonical reference output is itself). Then $\text{cl}_0(S_0) = \bigcup_{t\geq 0} g_L^t(S_0) = \{0,1,\ldots,2^L - 1\}$ already includes every state reachable along the orbit. Since $Q_L = g_L$, the descendant set satisfies $\mathcal{R}^{\text{rec}}_{Q_L}(S_0) = \text{cl}_0(S_0)$. Not strict containment. The counter chain leaves $S_0$ but does not exceed $\text{cl}_0(S_0)$; cumulative generation correctly fails.

#### Bounded-heredity corollary

The capacity and generation theorems give the bounded-heredity classification of substrates failing a strict subset of R1–R4.

**Corollary 41** (Bounded heredity, mode-keyed). *Within the (R5) scope, substrates failing a strict subset of R1–R4 exhibit heredity bounded according to which condition fails:*

1. *Failure of R1 at the per-site level (*$|\Sigma| = 1$*):* $C_S(L) = 0$. *No heritable distinguishability per site.*
2. *Failure of R1 at the macro level via a finite attractor repertoire of size* $K$*, with no extensible recognition substructure:* $C_S(L) \leq \log_2 K$*, length-independent. Mode 4: prion strains, amyloid attractors, DRT1 filament conformer.*
3. *Failure of R2 via a finite recurrent phase repertoire of* $N$ *states:* $C_S(L) \leq \log_2 N$*, regardless of product length. Mode 3: Drt3b at* $N = 2$ *gives* $C_S(L) \leq 1$ *bit.*

4. *Failure of R2 via finite module count $N$ with per-module alphabet of size $k$: $C_S(L) \leq N\log_2 k$. Mode 5: NRPS, polyketide synthases.*

5. *Failure of R3: substrate-level heredity is bounded by the information transmitted in a single application of $\mathcal{O}$; recursion is parasitic on a separate substrate–operator pair that itself satisfies R3. Mode 2 (translation: protein not ribosome-readable; recursion parasitic on Mode 1 acting on the gene encoding the ribosome). Mode 6 (cortex: surface does not autonomously self-copy; recursion parasitic on Mode 1 acting on the genome).*

6. *Failure of R4: substrate has heredity capacity but cannot generate novelty from a finite initial repertoire. Heredity is bounded by the founding-population repertoire.*

The corollary is a direct consequence of the upper bound in Proposition 29 together with the contrapositive structure of Theorems 36–39: each condition failure picks off a specific multiplicative or additive term in the bound. The biological assignment to Modes 1–6 follows the main paper's taxonomy table.

#### Finite-population proposition

The bounded-heredity corollary is a substrate-level statement (the heredity capacity ceiling at the kernel level). The corresponding population-level statement — what happens when a substrate–operator pair with a given failure pattern hosts a finite population under selection — requires careful scoping. The universal-cap formulation ("mechanisms failing R3 or R4 are structurally capped at $F_k^* < F_{\max}$ under any population dynamics") is too strong, because a wholesale-redraw mechanism with full support over $\mathcal{X}_L$ can sample any configuration by chance with infinite time. The proper finite-horizon, finite-population form follows.

**Proposition 42** (Finite-horizon plateau under nonlocal or absent drift). *Let a population of size $K$ evolve under selection sharpness $\beta$ toward a target phenotype on a tested-class landscape (single-target Hamming distance, $L$-dimensional alphabet, static), over a horizon $T < \infty$. Then:*

1. *Mechanisms whose substrate–operator pair fails R4 at the individual level (exact-copy systems: M1 lineage-level fixation, M3 individual-level exact copy) cannot exceed the maximum fitness $F_0^*$ present in the founding lineage repertoire. Mean fitness at horizon $T$ is bounded by $F_0^*$ regardless of $K$, $\beta$, or $T$.*

2. *Mechanisms whose substrate–operator pair has nonlocal R4 (wholesale redraw: M2 at rate $r > 0$) sample novelty with full support over $\mathcal{X}_L$ but do not accumulate local improvements. Under the tested update rules, the probability of hitting the target $X^*$ within horizon $T$ is bounded by $P(\tau \leq T) \; \leq \; 1 - \exp\left[-\frac{rKT}{|\mathcal{X}_L|}\right]$, with expected hitting time $\mathbb{E}[\tau] \sim |\mathcal{X}_L|/(rK)$. Since $|\mathcal{X}_L|$ scales as $|\mathcal{A}|^L$, the expected waiting time scales exponentially in $L$, and is negligible at biological scales for any fixed $K$, $r$, and biologically plausible $T$.*

3. *Mechanisms whose substrate–operator pair satisfies R3 and R4 with individual-level local drift (M4) are not structurally barred from cumulative climbing toward $F_{\max}$. Whether climbing actually occurs depends on landscape topology, finite-population drift, mutation load, neutral-network connectivity, and other conditions outside the framework's scope.*

The proposition is empirically tested in the main paper's Test H3–H5 sweep. The tested class ($L_{\text{target}} \in \{16,32,64,128\}$, $K = 400$, $\beta \in \{2,5,10,20\}$, 30 replicates per cell) confirms each item: M0 (no recursion) plateaus at chance (0.250); M1 and M3 (R4 fails) plateau near $F_0^* \approx 0.47–0.50$; M2 at sweet-spot $r^* = 0.10$ plateaus at $\sim 0.53$; M4 reaches $\sim 0.97$. The proposition's positive direction (M4 not structurally barred) is bounded: the simulation shows climbing on the tested landscape; it does not establish climbing on arbitrary landscapes.

#### Synthesis and recovery of prior formulations

The capacity and generation theorems clarify how prior formulations of the inheritance question fit within the framework. The correspondence in the table below is *conceptual* rather than mathematical equivalence: some entries are tight (Szathmáry's unlimited heredity is exactly $C_S(L) = \Omega(L)$ once the recursive-capacity definition is in place; Hull's replicator ontology is exactly R3 in its operational content), while others are interpretive in the sense that they identify which of the framework's conditions the prior formulation is addressing without claiming the two are formally equivalent statements (this is the appropriate reading of Pross's dynamic kinetic stability and Walker–Davies's top-down causation).

| Prior formulation | Conceptual correspondence | Reference |
|---|---|---|
| Schrödinger's aperiodic crystal | R1 + R2 (extensible recognition) | (Schrödinger 1944) |
| Szathmáry's unlimited heredity | $C_S(L) = \Omega(L)$ (capacity theorem) | (Szathmáry 2000) |
| Eigen's error threshold | Dynamical bound on $Q_L$ given R4, R5 | (Eigen 1971; Eigen et al. 1989) |
| Wagner's neutral networks | Topological condition on R4 drift in $\Sigma$ | (Wagner 2005, 2011) |
| Pross's dynamic kinetic stability | Thermodynamic-driving condition on R4 | (Pross 2011) |
| Hull's replicator ontology | R3 (substrate reads its own product) | (Hull 1980) |
| Maynard Smith's units of selection | R3 + R4 jointly | (Maynard Smith 2000) |
| Hofmeyr's $(F, A)$-systems | R3 in categorical form | (Hofmeyr 2017) |

| Prior formulation | Conceptual correspondence | Reference |
|---|---|---|
| Vasas et al. GARD refutation | Capacity-R2 failure for composition substrates | (Vasas et al. 2010) |
| Walker–Davies top-down causation | R3 in causal-flow form | (Walker and Davies 2013) |

The substantive contribution beyond these prior formulations is twofold: (i) the separation of *capacity* (R1, R2, R3) from *generation* (R4), which makes precise why a perfect copier transmits but does not innovate, and (ii) the explicit scope-defining role of R5, which marks the boundary between intrinsic catalytic chemistry and externally optimized search procedures.

Within this synthesis: Schrödinger's aperiodic-crystal intuition is precisely R1 (alphabet entropy) and R2 (extensible recognition) jointly; Szathmáry's unlimited heredity is exactly the capacity-theorem conclusion $C_S(L) = \Omega(L)$; Eigen's error threshold is a dynamical bound *on* $Q_L$ that becomes well-defined only once R4 holds and (R5) places the kernel in the intrinsic-physicochemistry domain; Wagner's neutral-network requirement is the topological structure of drift that R4 produces; Pross's dynamic kinetic stability supplies the thermodynamic driving under which R4 is sustainable; Hull's replicator ontology requires R3 (the substrate's product must be itself a substrate); Hofmeyr's $(F, A)$-systems give the categorical formulation of R3; Vasas et al.'s GARD refutation independently establishes capacity-theorem R2 failure for composition-vector substrates via a dynamical-fidelity route; Walker–Davies top-down causation supplies the causal-flow content of R3 in the information-substrate-to-phenotype direction.

The framework's value lies in the unification: each prior formulation addresses a particular condition or pair of conditions; the capacity/generation split shows precisely how the conditions combine and which biological chemistries fail which.

Cover, T. M., and J. A. Thomas. 2006. *Elements of Information Theory*. 2nd ed. Wiley-Interscience.

Eigen, M. 1971. "Selforganization of Matter and the Evolution of Biological Macromolecules." *Naturwissenschaften* 58: 465–523. https://doi.org/10.1007/BF00623322.

Eigen, M., J. McCaskill, and P. Schuster. 1989. "The Molecular Quasi-Species." *Advances in Chemical Physics* 75: 149–263.

Hofmeyr, J.-H. S. 2017. "Basic Biological Anticipation." In *Handbook of Anticipation*, edited by R. Poli. Springer.

Hull, D. L. 1980. "Individuality and Selection." *Annual Review of Ecology and Systematics* 11: 311–32. https://doi.org/10.1146/annurev.es.11.110180.001523.

Maynard Smith, J. 2000. “The Concept of Information in Biology.” *Philosophy of Science* 67: 177–94. https://doi.org/10.1086/392768.

Pross, A. 2011. “Toward a General Theory of Evolution: Extending Darwinian Theory to Inanimate Matter.” *Journal of Systems Chemistry* 2: 1. https://doi.org/10.1186/1759-2208-2-1.

Schrödinger, E. 1944. *What Is Life? The Physical Aspect of the Living Cell*. Cambridge University Press.

Szathmáry, E. 2000. “The Evolution of Replicators.” *Philosophical Transactions of the Royal Society B* 355: 1669–76. https://doi.org/10.1098/rstb.2000.0730.

Vasas, V., E. Szathmáry, and M. Santos. 2010. “Lack of Evolvability in Self-Sustaining Autocatalytic Networks Constraints Metabolism-First Scenarios for the Origin of Life.” *Proceedings of the National Academy of Sciences* 107: 1470–75. https://doi.org/10.1073/pnas.0912628107.

Wagner, A. 2005. *Robustness and Evolvability in Living Systems*. Princeton University Press.

Wagner, A. 2011. *The Origins of Evolutionary Innovations: A Theory of Transformative Change in Living Systems*. Oxford University Press.

Walker, S. I., and P. C. W. Davies. 2013. “The Algorithmic Origins of Life.” *Journal of the Royal Society Interface* 10 (79): 20120869. https://doi.org/10.1098/rsif.2012.0869.
